# Designing a reoriented H5 hemagglutinin to generate stem-directed, heterosubtypic immunity to influenza A viruses

**DOI:** 10.64898/2026.10.23.758699

**Authors:** Nicholas A. Miller, Tong Wang, Jung Ho Choi, Siyun Liang, Alena Yang, Peter J. Halfmann, Duo Xu

## Abstract

Human infections with the highly pathogenic avian influenza viruses highlight the risk of novel pandemics, against which seasonal vaccines provide limited cross-protection. Whereas antibody responses to current vaccine candidates are typically biased toward the immunodominant and variable head region of hemagglutinin (HA), the stem region of HA is highly conserved and targeted by broadly neutralizing antibodies. Here we describe a reoriented H5 HA (reoH5HA_Phe_, based on A/Vietnam/1203/2004) displayed on alum as a vaccine candidate to elicit stem-directed antibody responses and confer cross-protective immunity. We show that a single immunization with reoH5HA_Phe_ protected naïve mice against heterologous H5N1 lethal challenge (A/Texas/37/2024), and boosting with reoH5HA_Phe_ conferred complete heterologous protection in mice pre-immunized with seasonal vaccines. The stem-directed response elicited by reoH5HA_Phe_ also protected against heterosubtypic H1N1 challenge in both naïve and pre-immune mice. Electron microscopy polyclonal epitope mapping revealed that reoH5HA_Phe_ elicited antibodies targeting the central stem and anchor epitopes on heterologous H5 and H1 HAs, alongside antibodies against the (H5) vestigial esterase or (H1) lateral patch epitopes. Despite limited cross-neutralizing activity, increased stem-directed responses with balanced IgG1 and IgG2 subclasses reduced viral replication and improved survival. Our results highlight antigen reorientation as an effective approach for designing variant-resilient vaccines and demonstrate the potential of reoH5HA_Phe_ as an H5 pandemic influenza vaccine candidate that confers protective immunity against both emerging and seasonal influenza viruses.

**Significance Statement:** The global spread and human infections of avian H5N1 influenza viruses raise concerns of novel pandemics, against which humans lack pre-existing immunity. Whereas current vaccine strategies, including mRNA vaccines, mostly protect against sequence-matched viruses, pandemic preparedness requires vaccine design that resists antigenic variation and viral evolution. Here we employ molecular control of antigen orientation to focus antibody responses toward evolutionarily conserved sites of influenza hemagglutinin. We designed a reoriented H5 hemagglutinin based on the historical sequence (A/Vietnam/1203/2004) and demonstrated robust protection against the contemporary virus (A/Texas/37/2024). Vaccinations with reoriented H5 hemagglutinin also conferred protection against historical and contemporary H1N1 viruses. Our approach has potential to provide protection against emerging, pandemic-potential viruses, while simultaneously boosting immunity to seasonal influenza viruses.

## Introduction

Human infections with the highly pathogenic avian influenza (HPAI) H5N1 viruses are historically associated with severe disease and high fatality(1, 2). Recent studies have identified genetic mutations in H5 hemagglutinin (HA) that favor viral adaptation to human infection(3, 4), highlighting the risk of influenza pandemics, against which seasonal vaccine-induced immunity is unlikely to provide cross-protection(5). The recent cattle-to-human transmission of the H5N1 clade 2.3.4.4b virus underscores such risks and the urgent need to develop vaccine candidates that protect against antigenically distinct and emerging influenza viruses(6, 7).

Vaccine candidates based on HPAI H5N1 viruses have been developed, and they induce neutralizing antibody responses and protection against homologous lethal challenge(8–16). Nonetheless, similar to seasonal vaccines, they primarily elicit antibodies to the immunodominant and variable head region of HA (HA-head) that undergoes antigenic drift(17, 18). As a result, vaccines based on current strains may require updates to antigenically match emerging variants, leading to a variant-chasing strategy for pandemic responses(19, 20). Developing broad immunity against antigenically diverse influenza viruses therefore requires vaccine candidates to overcome HA-head immunodominance and focus immune responses (or immunofocus) on more conserved antigenic regions(21). Compared with HA-head, the stem region of HA (HA-stem) is immunosubdominant but highly conserved and targeted by broadly neutralizing antibodies (bnAbs)(22–26). Prophylactic treatment with stem-directed bnAbs protects nonhuman primates against HPAI H5N1 lethal challenge(27), and stem-specific antibodies correlate with reduced infection and disease in humans(28), suggesting that stem-based vaccine strategies can generate broadly protective immunity.

Current stem-based vaccine strategies include display of heterologous or inverted HAs(29–32), chimeric HAs(33, 34), epitope masking of HA-head(35–37), and stem-only immunogens(38, 39). These strategies have successfully expanded the breadth of antibody responses but often require multi-component assembly, repeated and heterologous immunization regimens, or iterative protein engineering to shield or preserve conformational epitopes, respectively. We recently introduced antigen reorientation as an orthogonal approach to designing epitope-focused vaccines by presenting antigens on aluminum hydroxide (alum) adjuvants in a predefined orientation(40). Instead of epitope masking or protein dissection, our approach uses full-length antigen proteins to preserve their native conformation and site-specific insertion of aspartate residues (oligoD) to control antigen orientation on alum. Reorientation of HA on alum enhances exposure of native stem epitopes while sterically occluding HA-head to reduce its immunodominance. We previously showed that a reoriented H2 HA enhanced elicitation of stem-directed antibodies, although it remains unknown whether this response is functional or protective *in vivo*, particularly against antigenically distinct or emerging influenza viruses.

Because the genetic sequence of future pandemic viruses is unpredictable, we establish a historical-to-contemporary scenario to test whether antigen reorientation confers broad immunity and reduces the need for antigenic matching. Here we investigate whether a reoriented H5 HA (based on A/Vietnam/1203/2004 or VN/04 with a phenylalanine substitution on HA-stem(41), referred to as reoH5HA_Phe_) elicits a stem-directed, protective response against emerging H5N1 viruses (A/Texas/37/2024) while simultaneously enhancing immunity against seasonal influenza strains. We show that compared to wild-type H5 HA_Phe_, reoH5HA_Phe_ on alum elicited a more stem-directed response that broadly cross-reacted with different group 1 influenza A HAs. In naïve mice, a single immunization with reoH5HA_Phe_ protected 50% or more of mice against heterologous lethal challenge with the recent HPAI H5N1 clade 2.3.4.4b virus (A/Texas/37/2024, TX/24)(42), whereas H5 HA_Phe_ did not. In mice with pre-existing immunity established by seasonal vaccines, a single booster immunization with reoH5HA_Phe_ provided complete heterologous protection against TX/24, whereas boosting with wild-type H5 HA_Phe_ or the seasonal vaccine conferred minimal or no protection, respectively. Negative-stain electron microscopy polyclonal epitope mapping (nsEMPEM) showed that only reoH5HA_Phe_ elicited cross-reactive antibodies targeting distinct stem epitopes on the heterologous H5 HA (TX/24).

Additionally, immunizations with reoH5HA_Phe_ induced H1-stem-reactive antibodies and protected against H1N1 viruses in both naïve and pre-immune mice. nsEMPEM analysis revealed that only reoH5HA_Phe_ elicited cross-reactive antibodies toward the lateral patch and stem epitopes on the intact H1 HA trimer. Although antisera lacked cross-neutralizing activity against heterologous H5N1 and H1N1 viruses, high levels of stem-directed antibodies with balanced IgG1 and IgG2 subclasses led to reduced viral replication in respiratory tissues and improved *in vivo* protection. Our results demonstrate antigen reorientation as an effective approach for designing epitope-focused vaccines against antigenically diverse viruses and showcase the potential of reoH5HA_Phe_ as an H5 pandemic influenza vaccine candidate.

## Results

### Design and characterization of reoH5HA

Based on the reoriented H2 HA developed previously(40), we aligned the structure of H5 and H2 HAs (**Fig. S1a**) and identified the oligoD insertion site after residue N164 (H3-numbering) to generate reoH5HA (**Fig. 1a**). We also introduced a phenylalanine substitution on HA-stem (Phe45_HA2_, **Fig. 1b**), since this bulky residue from H2 HAs was shown to expand the breadth of human B cell and antibody responses(41). H5 HA, reoH5HA, and their Phe45_HA2_ variants (H5 HA_Phe_ and reoH5HA_Phe_) were expressed in Expi293F cells and purified to homogeneity (**Fig. 1c, Fig. S1b,c**). Although oligoD insertion decreased the thermal melting temperature (T_m_) of H5 HA and H5 HA_Phe_ by 2 to 3 °C in solution, T_m_ of reoH5HA and reoH5HA_Phe_ was higher than their wild-type counterparts in the presence of alum, suggesting increased thermal stability *via* alum-binding (**Fig. 1d, Fig. S1d**). To assess antigenicity, we used HA-specific monoclonal antibodies (mAbs) that recognize the vestigial esterase (VE) or stem epitopes (**Fig. S1e**) and examined their binding to wild-type and modified H5 HAs by biolayer interferometry (BLI, **Fig. 1e, Fig. S1f**). All mAbs showed similar high-affinity binding, except for CR9114 with reduced binding to H5 HA_Phe_ and reoH5HA_Phe_, since the Phe45_HA2_ substitution is known to sterically clash with CR9114 paratopes(43).

**Figure 1.**
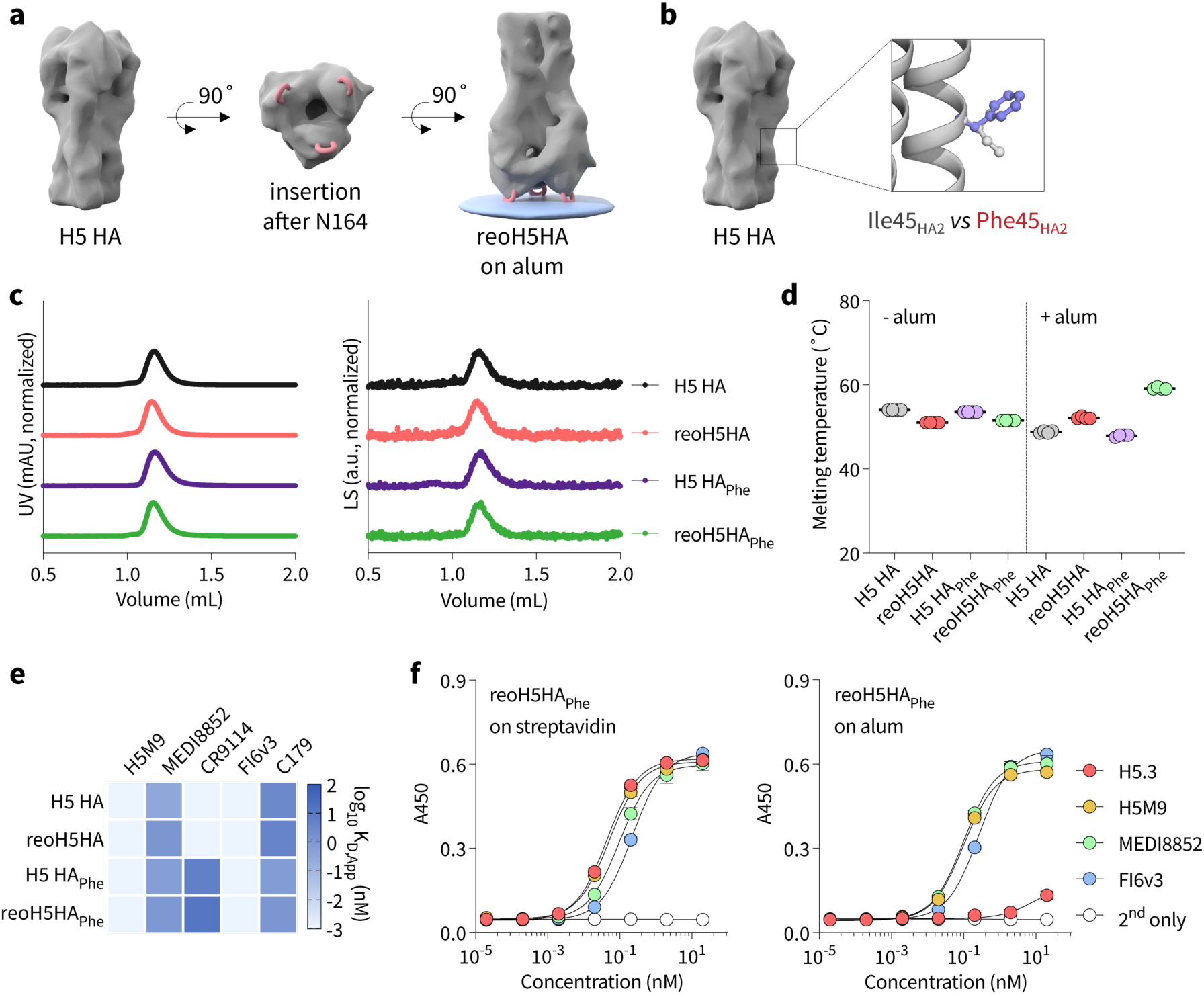
Design of reoH5HA with Phe45_HA2_ substitution. **a**, Insertion of oligoD into the head region of H5 HA (VN/04) after residue N164 (H3 numbering) enabled its binding to alum in an “upside-down” orientation. **b**, Comparison of the wild-type Ile45_HA2_ residue and modeled Phe45_HA2_ residue on HA-stem or the HA2 subunit. **c**, Size-exclusion chromatography coupled with multi-angle light scattering (SEC-MALS) analysis of H5 HA, H5 HA_Phe_, reoH5HA, and reoH5HA_Phe_ on a Superdex 200 column. Normalized UV and LS indicate absorbance at 280 nm and light scattering signals, respectively. **d**, Thermal melting temperature (T_m_) of wild-type and modified H5 HAs in the absence or presence of alum. Data are presented as mean ± s.d. (*n =* 4 technical replicates). **e**, BLI binding analysis of HA-specific mAb to wild-type or modified H5 HA. Log_10_-transformed apparent binding affinity (K_D,App_) of each mAb was determined based on **fig. S1f**. **f**, Binding of head-(H5.3), VE-(H5M9) or stem-directed mAbs (MEDI8852 or FI6v3) to reoH5HA_Phe_ on streptavidin-or alum-based ELISA plates. Data are presented as mean ± s.d. (*n =* 4 technical replicates).

We next assessed the orientation of reoH5HA and reoH5HA_Phe_ on alum using streptavidin- and alum-based enzyme-linked immunosorbent assays (ELISAs) with head-, VE-, and stem-directed mAbs (**Fig. S1g**). When wild-type and modified H5 HA proteins were C-terminally biotinylated and captured on streptavidin-coated plates, all mAbs exhibited similar high-affinity binding, indicating that these epitopes were preserved and accessible (**Fig. 1f, Fig. S1h**). H5 HA and H5 HA_Phe_ poorly adsorbed to alum without oligoD modification, with negligible mAb binding in alum-based ELISAs (**Fig. S1i**). By comparison, when reoH5HA and reoH5HA_Phe_ were adsorbed on alum-coated plates, all mAbs retained their high-affinity binding except the head-directed H5.3 mAb (**Fig. 1f, Fig. S1h**), suggesting steric occlusion of its epitope on HA-head. Similar to the previous study(40), comparison of streptavidin- and alum-based ELISAs validated the upside-down orientation of reoH5HA and reoH5HA_Phe_ on alum.

### Reorientation of H5 HA enhances stem-directed antibody responses

We immunized mice with H5 HA_Phe_ or reoH5HA_Phe_ adjuvanted with alum and CpG oligodeoxynucleotide *via* intramuscular injections to assess their immunogenicity (**Fig. 2a**). A prime-boost regimen with H5 HA_Phe_ or reoH5HA_Phe_ induced similar H5 HA-specific IgG responses (**Fig. 2b**) and neutralizing activity against the homologous H5N1 pseudovirus (VN/04) (**Fig. 2c, Fig. S2a**). Despite comparable H5 HA-specific responses, reoH5HA_Phe_ elicited significantly higher IgG titers against HA-stem, as measured with H5 stabilized stem (H5-SS, **Fig. 2d, Fig. S2b**). Increased stem-directed responses translated into broad cross-reactivity with different group 1 HAs, with antisera raised against reoH5HA_Phe_ showing higher and more consistent binding than those raised against H5 HA_Phe_ (**Fig. 2e**). Antisera from both groups also competed binding with stem-(MEDI8852, **Fig. S2c**) and anchor-directed bnAbs (222-1C06, **Fig. S2d**)(26, 44), with greater competition from the reoH5HA_Phe_ group. Nonetheless, neither group developed strong cross-neutralizing activity against the heterologous H5N1 pseudovirus (TX/24) (**Fig. S2e,f**). Similar to previous studies(40), use of a high dose of alum (150 µg) induced a Th2-biased response with substantially higher titers of H5 HA-specific IgG1 than IgG2 subclasses (**Fig. S2g**). This bias was more pronounced for H5-SS-specific responses, in which binding was almost exclusively contributed by IgG1 from both immunogens (**Fig. S2h**).

**Figure 2.**
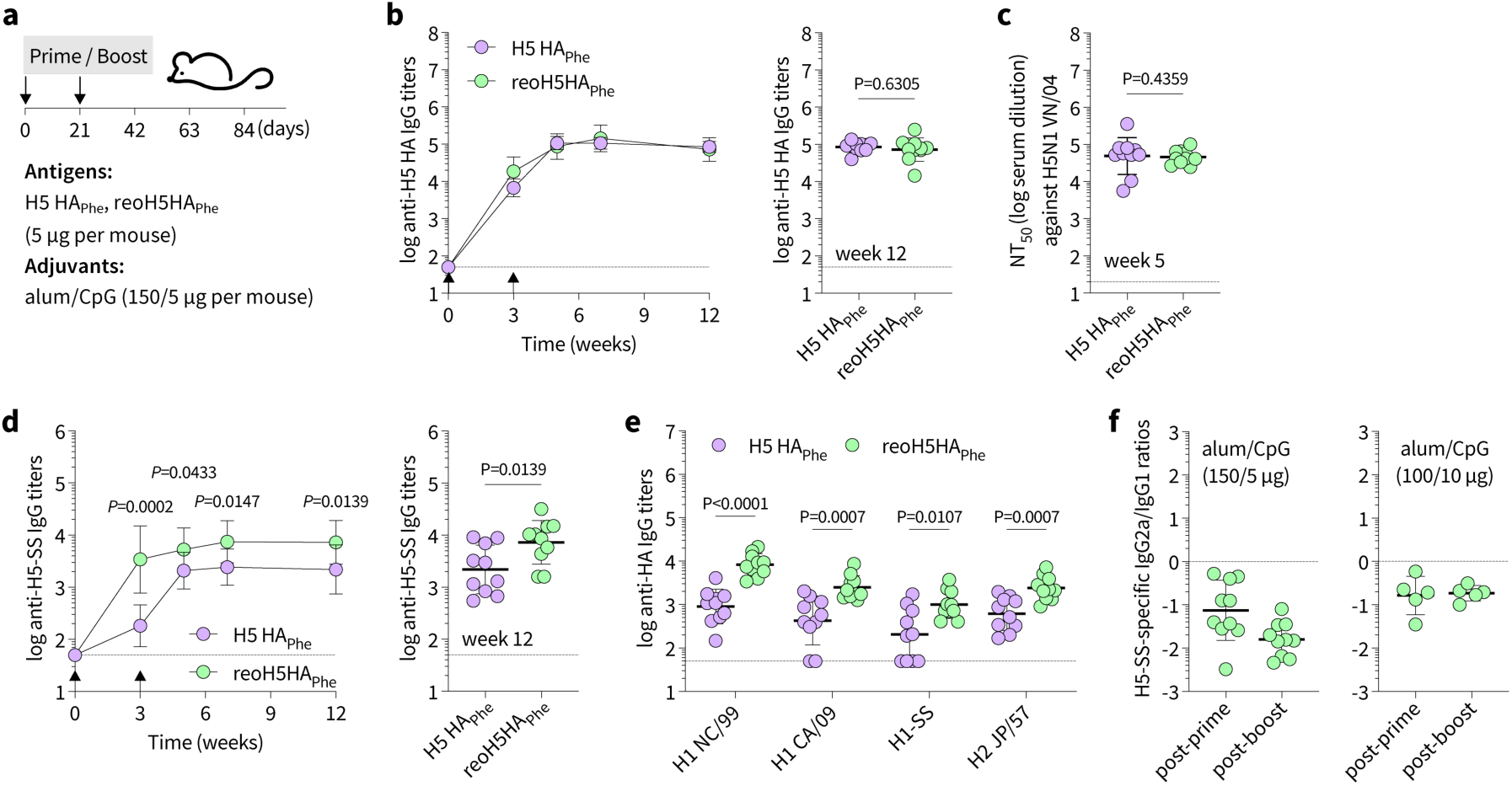
Immunization with reoH5HA_Phe_ enhances stem-directed antibody responses. **a**, A prime-boost immunization with H5 HA_Phe_ or reoH5HA_Phe_ (5 µg per mouse) adjuvanted with alum/CpG (150/5 µg per mouse) *via* intramuscular injection in female BALB/c mice (*n =* 10 per group). **b**, Serum H5 HA-specific IgG titers over time. Endpoint (week 12) IgG titers are plotted on the right for comparison. **c**, Comparison of neutralization titers (NT_50_) against the homologous H5N1 pseudovirus (VN/04) by week 5 antisera. **d**, Serum H5-SS-specific IgG titers over time. Endpoint (week 12) antibody titers are plotted on the right for comparison. **e**, Cross-reactive binding of group 1 HAs (H1 NC/99, H1 CA/09, H1-SS, and H2 JP/57) by week 7 antisera. **f**, Ratios of H5-SS-specific IgG2a/IgG1 subclasses following prime-boost immunizations with different doses of alum/CpG adjuvants (150/5 µg *vs.* 100/10 µg per mouse). Arrows indicate prime and boost immunizations in **b,d**. Data are presented as geometric mean ± s.d. of the log_10_- transformed values in **b-f**. Dashed lines indicate the limit of quantification in **b-e** and the ratio of 1.0 in **f**. Comparison of two groups was performed using the two-tailed Mann-Whitney U-test. P values of 0.05 or less were considered significant.

### Stimulating a balanced, stem-directed response via adjuvant screening

Since protection by stem-directed, non-neutralizing antibodies is mainly driven by Fc-mediated effector functions and murine IgG2 subclasses correlate with stronger effector functions than murine IgG1(38, 45), we examined different alum-compatible adjuvant combinations to address the Th2-biased response while preserving reoH5HA_Phe_ binding to alum. Given that CpG contains phosphate groups that compete for alum binding with antigen, we titrated reoH5HA_Phe_/alum/CpG ratios to identify the permissible amount of CpG in the antigen-adjuvant mixture (**Fig. S3a**). In the absence of CpG, reoH5HA_Phe_ remained alum-bound at multiple reoH5HA_Phe_/alum ratios (1/10, 20, or 30, *w*/*w*) (**Fig. S3b**). Whereas low ratios of CpG (0.5, 1, or 2/1, CpG/reoH5HA_Phe_, *w*/*w*) had minimal effects on reoH5HA_Phe_ binding to alum, high ratios of CpG (4, 8, or 10/1, CpG/reoH5HA_Phe_, *w*/*w*) reduced the alum-bound fraction of reoH5HA_Phe_ (**Fig. S3c**). Based on this titration, we adjusted the reoH5HA_Phe_/alum/CpG ratio to 1/20/2 (*w*/*w*) and performed an *in vivo* adjuvant screening with alum/CpG and four additional alum-compatible adjuvants (**Fig. S3d**). Among all, use of alum/CpG or alum/monophosphoryl lipid A stimulated the highest H5 HA- and H5-SS-specific IgG responses (**Fig. S3e,f**), while alum/CpG induced a more balanced response with elevated IgG2 titers against H5-SS (**Fig. 2f, Fig. S3g**). Increased IgG2 levels also promoted binding to the mouse Fc gamma receptors (mFcγRIIIa and mFcγRIV, **Fig. S3h**), which are essential for Fc-mediated effector functions(46). Additional titration at a lower reoH5HA_Phe_/alum ratio (1/100, *w*/*w*) revealed that reoH5HA_Phe_ remained completely alum-bound across a wide range of CpG ratios (1/1 to 1/25, CpG/reoH5HA_Phe_, *w*/*w*, **Fig. S3i,j**), providing a broad window for the antigen-adjuvant mixture without affecting antigen reorientation.

### A single immunization with reoH5HA_Phe_ protects against heterologous H5N1 lethal challenge

We next performed a single-dose immunization with H5 HA_Phe_ or reoH5HA_Phe_ formulated with alum/CpG at two antigen-adjuvant ratios (**Fig. 3a**). Both immunogens induced robust H5 HA-specific antibody responses, with the highest titers elicited by reoH5HA_Phe_ (1 µg) adjuvanted with alum and high-dose CpG (100 and 25 µg) (**Fig. 3b, Fig. S4a**). H5 HA-specific IgG titers correlated with neutralizing activity against the homologous H5N1 pseudovirus (**Fig. S4b**), in which reoH5HA_Phe_ with alum and high-dose CpG induced the highest neutralization titers (NT_50_, **Fig. 3c**). H5 HA_Phe_ (1 µg) adjuvanted with alum and high-dose CpG (100 and 25 µg) induced the lowest IgG titers and NT_50_, presumably due to poor alum binding and limited antigen availability post-vaccination. With low-dose CpG (10 µg), H5 HA_Phe_ and reoH5HA_Phe_ (5 µg) on alum (100 µg) induced similar H5 HA-specific IgG titers, whereas reoH5HA_Phe_ elicited almost tenfold higher H5-SS-specific responses after week 4 (**Fig. 3d, Fig. S4c**), indicating a more immunofocused response to HA-stem. With high-dose CpG (25 µg), reoH5HA_Phe_ on alum elicited approximately 30- and 50-fold higher H5 HA- and H5-SS-specific IgG titers than H5 HA_Phe_, respectively, with a higher fraction of antibodies directed toward HA-stem. Increased stem-directed responses enhanced cross-reactivity with HAs from HPAI H5N1 viruses (**Fig. 3e**), despite limited cross-neutralizing activity (**Fig. S4d,e**). To structurally characterize epitopes targeted by antisera, we performed nsEMPEM with the heterologous H5 HA (TX/24) and found that while H5 HA_Phe_ only induced cross-reactive antibodies against the VE epitope, reoH5HA_Phe_ elicited antibodies targeting the VE, central stem, and anchor epitopes (**Fig. 3f, Fig. S4f**). Use of optimized alum/CpG ratios also induced balanced, stem-directed IgG1 and IgG2 subclasses over time (**Fig. S4g-j**), where elevated IgG2 titers correlated with increased binding to mFcγRIIIa and mFcγRIV (**Fig. S4k**).

**Figure 3.**
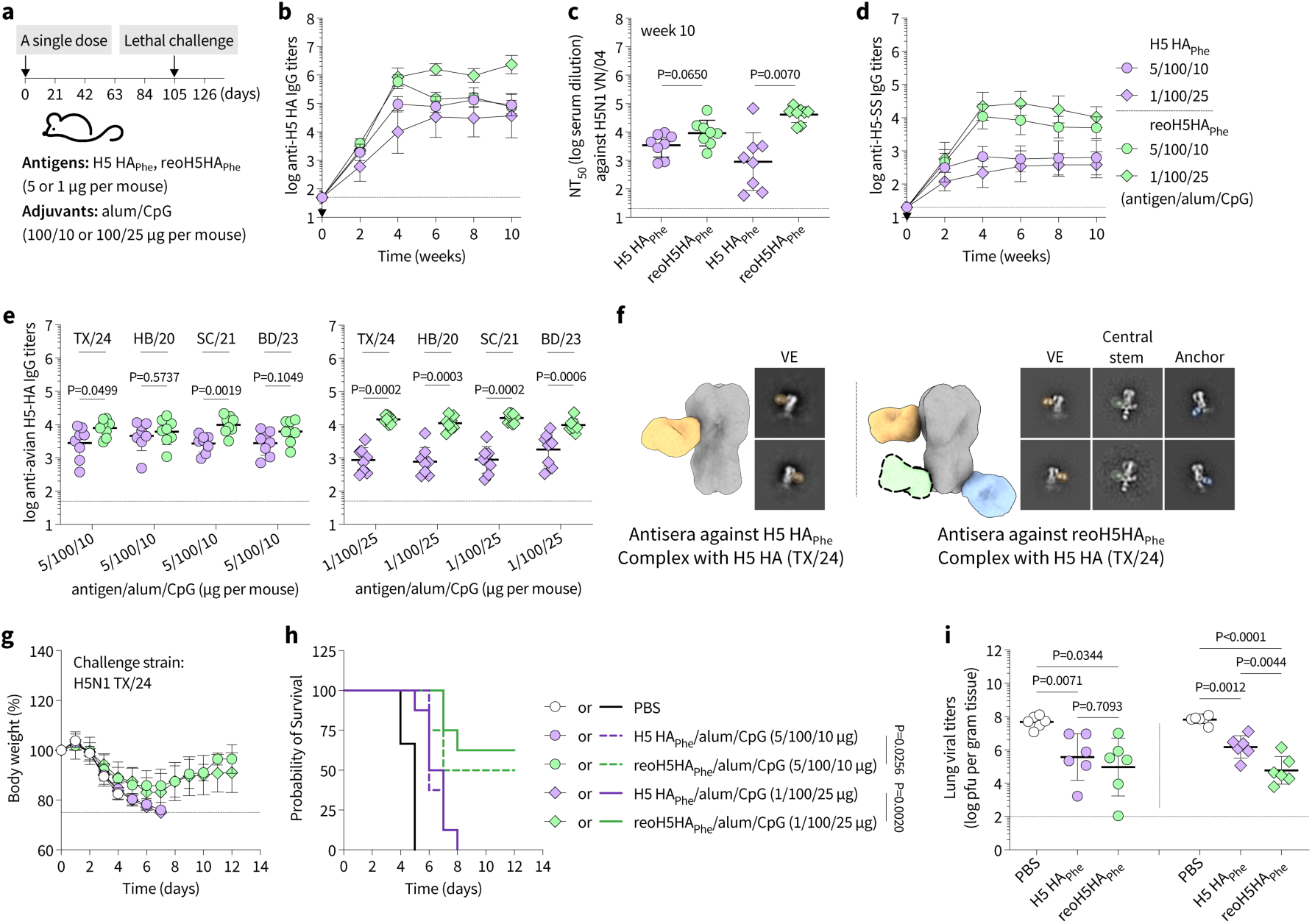
A single immunization with reoH5HA_Phe_ protects against heterologous H5N1 lethal challenge. **a,** A single-dose immunization with H5 HA_Phe_ or reoH5HA_Phe_ (5 or 1 µg per mouse) adjuvanted with alum/CpG (100/10 or 100/25 µg per mouse) *via* intramuscular injection in female BALB/c mice (*n =* 8 per group). **b**, Serum H5 HA-specific IgG titers over time. **c**, Comparison of NT_50_ against the homologous H5N1 pseudovirus (VN/04) by week 10 antisera. **d**, Serum H5-SS-specific IgG titers over time. **e**, Cross-reactive binding of HAs from HPAI H5N1 strains by week 10 antisera. **f**, nsEMPEM analyses for pooled antisera raised against H5 HA_Phe_ or reoH5HA_Phe_. 3D composite maps and representative 2D class averages generated from negative-stain reconstructions of polyclonal immune complexes with H5 HA (TX/24) are shown for each group, with dotted lines highlighting the likely position of the Fab population at low abundance. **g,** Body weight loss and **h**, Kaplan-Meier survival curves of mice after lethal challenge (10× mLD_50_) with a heterologous H5N1 virus (TX/24). Comparison of survival curves was performed using a pairwise log-rank test. **I**, Female BALB/c mice were immunized on day 0 with PBS, H5 HA_Phe_, or reoH5HA_Phe_ (5 or 1 µg per mouse) adjuvanted with alum/CpG (100/10 or 100/25 µg per mouse) and challenged with H5N1 virus (10× mLD_50_, TX/24) on day 105 (*n =* 6 per group). Lung viral titers were assessed three days post-infection. Comparison of viral titers was performed using lognormal one-way ANOVA followed by a Tukey’s post hoc test. Arrows indicate the single-dose immunization in **b,d**. Data are presented as geometric mean ± s.d. of the log_10_-transformed values in **b-e**,**i** or mean ± s.d. in **g**. Dashed lines indicate the limit of quantification in **b-e,i** or 25% weight loss in **g**. Comparison of two groups was performed using the two-tailed Mann-Whitney U-test in **c,e**. P values of 0.05 or less were plotted and considered significant.

Fourteen weeks after the single-dose immunization, we performed lethal challenge with the HPAI H5N1 clade 2.3.4.4b virus (TX/24, 10× mouse lethal dose 50 or 10× mLD_50_, **Fig. S4l**) and monitored body weight loss and survival (**Fig. 3g,h**). As the control, mice immunized with PBS rapidly lost weight and succumbed to infection within 4 to 5 days. Although H5 HA_Phe_ with alum/CpG elicited a robust antigen-specific response (**Fig. 3b,c**), it failed to protect against the lethal challenge. By contrast, a single immunization with reoH5HA_Phe_ and alum/CpG protected 50% or more mice, with high-dose CpG further improving protective efficacy. In a parallel study, we collected lung, nasal turbinate, and brain tissues from mice three days post-infection to assess viral replication. Although both immunogens reduced viral replication relative to the control group (**Fig. 3i, Fig. S4m,n**), reoH5HA_Phe_ further decreased viral titers in the respiratory tissues, correlating with improved survival.

In addition to the single-dose immunization, we examined a prime-boost regimen with H5 HA_Phe_ or reoH5HA_Phe_ adjuvanted with alum/CpG (**Fig. S5a**). Consistent with previous immunizations, reoH5HA_Phe_ elicited a substantially higher stem-directed response with both IgG1 and IgG2 subclasses than H5 HA_Phe_ (**Fig. S5b**). Despite differences in antibody titers, both H5 HA_Phe_ and reoH5HA_Phe_ conferred complete protection against lethal challenge with heterologous HPAI H5N1 viruses with distinct genotypes, TX/24 (B3.13 genotype, **Fig. S5c,d**) and A/Washington/239/2024 (D1.1 genotype, **Fig. S5e,f**).

### A booster immunization with reoH5HA_Phe_ confers complete heterologous protection

To simulate pre-existing immunity to influenza virus, we primed mice with a tetravalent seasonal influenza vaccine, Fluzone (FZ, including H1N1-A/Victoria/4897/2022 IVR-238, H3N2-A/Darwin/9/2021 SAN-010, B/Phuket/3073/2013, and B/Michigan/01/2021) (**Fig. 4a**). After the prime, all mice developed high titers of antigen-specific antibodies against HAs from the seasonal vaccine strains (**Fig. S6a**), although with limited stem-directed responses (**Fig. S6b,c**) and minimal cross-reactivity to H5 HAs (**Fig. S6d**). Four weeks post-prime, we boosted mice with PBS, FZ, H5 HA_Phe_, or reoH5HA_Phe_. Whereas FZ further elevated antibody responses to seasonal influenza HAs with minimal cross-reactivity to H5 HAs (**Fig. S6e,f**), H5 HA_Phe_ and reoH5HA_Phe_ barely boosted responses to seasonal strains but primarily induced H5 HA-specific antibodies with limited neutralizing activity against the homologous H5N1 virus (VN/04) (**Fig. S6g,h**). reoH5HA_Phe_ also induced a higher stem-specific response than H5 HA_Phe_ (**Fig. 4b, Fig. S6i**) with increased cross-reactivity to heterologous H5 HAs, including TX/24 (**Fig. 4c, Fig. S6j**), despite minimal cross-neutralization (**Fig. S6k**). Since only reoH5HA_Phe_ elicited appreciable cross-reactivity to H5 HA (TX/24) two weeks post-boost, we performed nsEMPEM only with antisera raised against reoH5HA_Phe_ and observed cross-reactive antibodies targeting the central stem epitope of H5 HA (TX/24, **Fig. 4d**). Six weeks post-boost, we performed lethal challenge with the heterologous H5N1 virus (TX/24, 10× mLD_50_) and monitored body weight loss and survival (**Fig. 4e,f**). While one or two doses of FZ immunization did not provide any protection, boosting with H5 HA_Phe_ protected 2/10 mice. By contrast, mice boosted with reoH5HA_Phe_ displayed reduced weight loss and all survived the challenge. In a parallel study, we assessed viral replication in lung, nasal turbinate, and brain tissues three days post-infection (**Fig. 4g**) and found that only reoH5HA_Phe_ significantly reduced viral loads in respiratory tissues (**Fig. S6l**), correlating with improved survival.

**Figure 4.**
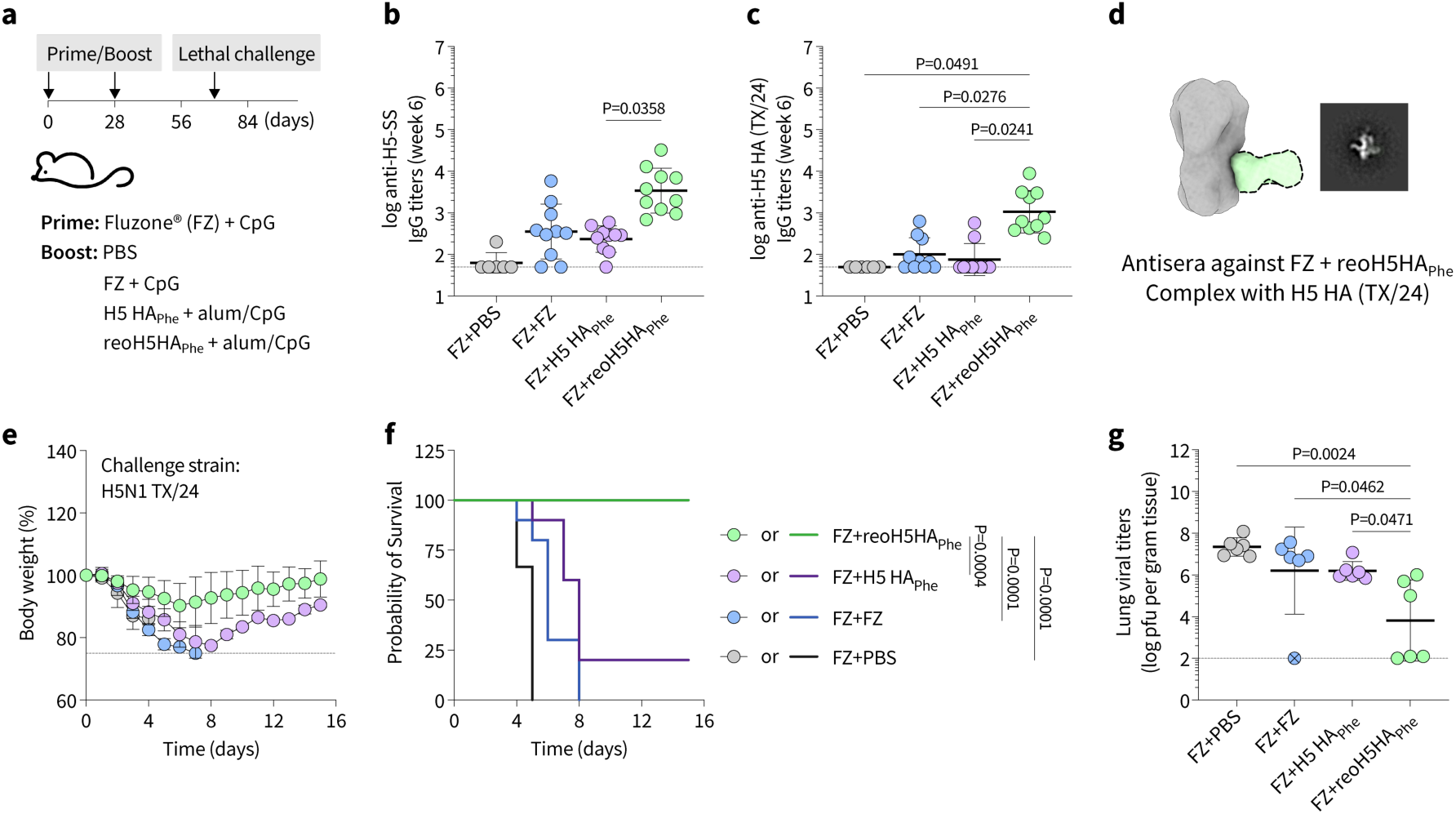
A single boost with reoH5HA_Phe_ confers complete protection against heterologous H5N1 lethal challenge in mice with pre-existing immunity. **a**, A prime-boost immunization *via* intramuscular injection in female BALB/c mice. All mice were primed with Fluzone (FZ, 20 µL per mouse) adjuvanted with CpG (10 µg per mouse) and then boosted with PBS (*n =* 6), FZ adjuvanted with CpG (*n =* 10), H5 HA_Phe_ (1 µg per mouse) adjuvanted with alum/CpG (100/25 µg per mouse, *n =* 10), or reoH5HA_Phe_ (1 µg per mouse) adjuvanted with alum/CpG (100/25 µg per mouse, *n =* 10). **b**, Serum H5-SS-specific and **c**, H5 HA (TX/24) cross-reactive IgG titers by week 6 antisera from all groups. **d**, nsEMPEM analyses for pooled antisera raised against the FZ + reoH5HA_Phe_ group. The 3D composite map and representative 2D class average generated from negative-stain reconstruction of the polyclonal immune complex with H5 HA (TX/24) are shown, with dotted lines highlighting the likely position of the Fab population at low abundance. **e**, Body weight loss and **f**, Kaplan-Meier survival curves of mice after lethal challenge (10× mLD_50_) with a heterologous H5N1 virus (TX/24). Comparison of survival curves was performed using a pairwise log-rank test. **g**, Female BALB/c mice were immunized with FZ + PBS, FZ + FZ, FZ + H5 HA_Phe_, or FZ + reoH5HA_Phe_ to assess viral replication post-lethal challenge (*n =* 6 per group). Lung viral titers were assessed three days post-infection with H5N1 virus (10× mLD_50_, TX/24). Comparison of viral titers was performed using lognormal one-way ANOVA followed by a Tukey’s post hoc test. The crossed circle in **g** indicates unsuccessful viral infection. Data are presented as geometric mean ± s.d. of the log_10_-transformed values in **b,c,g** or mean ± s.d. in **e.** Dashed lines indicate the limit of quantification in **b,c,g** or 25% weight loss in **e**. Comparison of IgG titers and NT_50_ from multiple groups was performed using one-way ANOVA followed by a Tukey’s post hoc test. P values of 0.05 or less were plotted and considered significant.

### reoH5HA_Phe_ elicits cross-reactive antibodies to H1-stem and protects against H1N1 challenge

We next examined whether the stem-directed response elicited by reoH5HA_Phe_ protected against heterosubtypic influenza viruses (*e.g.*, H1N1), in addition to H5N1 subtypes. Considering the antigenic difference between H5 and H1 HAs (**Fig. S7a**), we immunized mice three times with H5 HA_Phe_ or reoH5HA_Phe_ adjuvanted with alum/CpG (**Fig. 5a**). Both immunogens induced high titers of H5 HA-specific IgG with homologous neutralizing activity (**Fig. 5b, Fig. S7b,c**). Despite similar H5 HA-specific responses, reoH5HA_Phe_ elicited significantly higher IgG titers against both H5-SS and H1-SS than H5 HA_Phe_ over time (**Fig. 5c,d**), with balanced IgG1 and IgG2 subclasses (**Fig. S7d,e**), suggesting a more stem-directed response. Whereas H5 HA_Phe_ only induced cross-reactive antibodies bound to HA protomers under the nsEMPEM analysis, reoH5HA_Phe_ elicited antibodies targeting the lateral patch, central stem, and anchor epitopes on the intact H1 HA trimer (**Fig. 5e**). Four weeks after the third immunization, we performed lethal challenge with the 2009 H1N1 pandemic virus (H1N1pdm09 or A/Netherlands/602/2009, 10× or 15× mLD_50_) and monitored body weight loss and survival (**Fig. 5f,g, Fig. S7f,g**). At both challenge doses, PBS-immunized mice rapidly lost weight and succumbed to infection within 6 to 8 days. Whereas immunizations with H5 HA_Phe_ protected only 1/8 mice at both challenge doses, reoH5HA_Phe_ reduced weight loss and protected 6/8 mice at 10× mLD_50_ and 5/8 mice at 15× mLD_50_, despite negligible cross-neutralizing activity (**Fig. S7h**). As the positive control, mice immunized with H1 HA (A/California/07/2009) developed neutralizing antibody responses and all survived (**Fig. S7i,j**).

**Figure 5.**
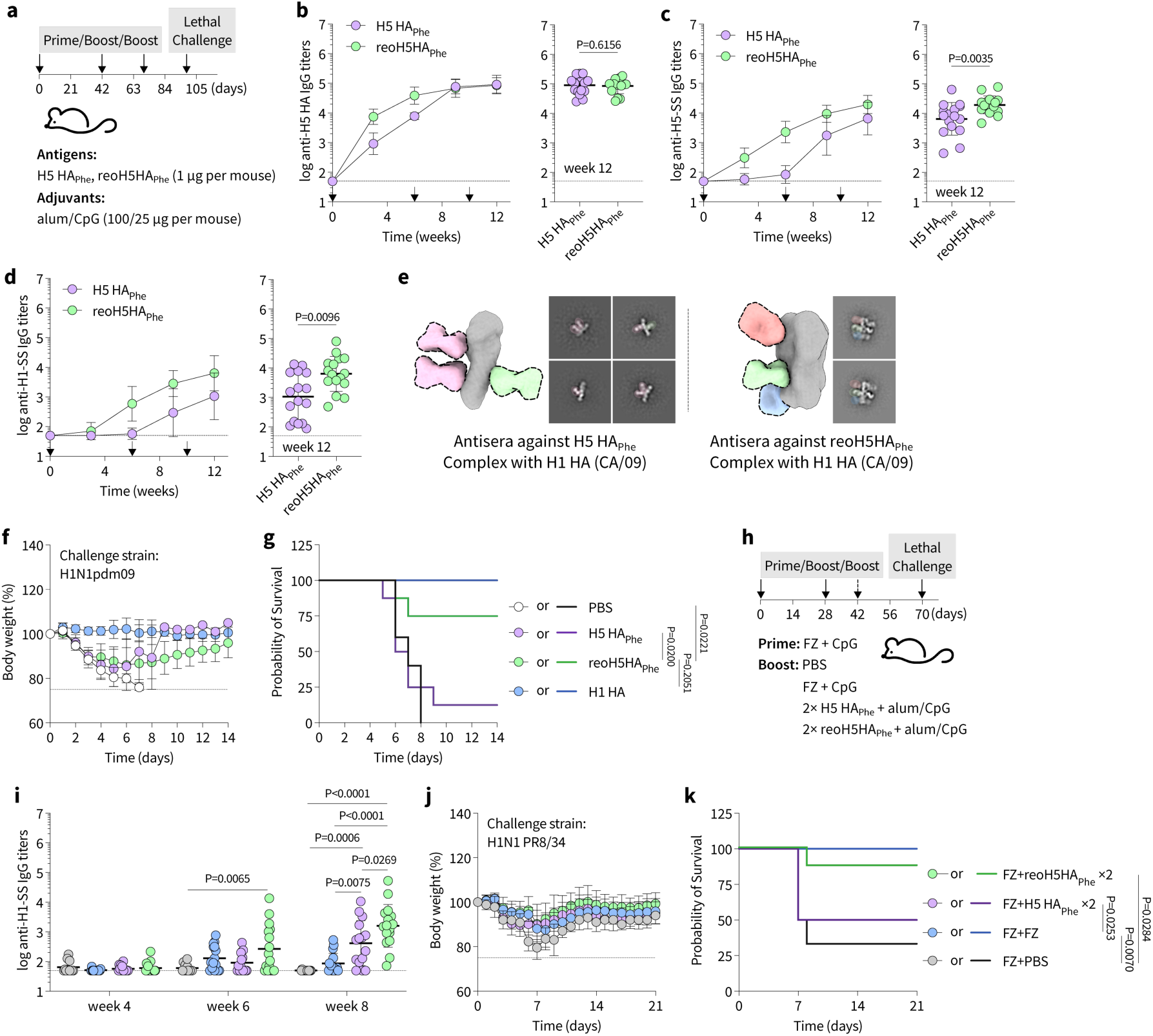
Immunizations with reoH5HA_Phe_ confer heterosubtypic immunity against H1N1 viruses. **a**, A three-dose immunization with H5 HA_Phe_ or reoH5HA_Phe_ (1 µg per mouse) adjuvanted with alum/CpG (100/25 µg per mouse) *via* intramuscular injection in female BALB/c mice (*n =* 16 per group). **b**, Serum H5 HA-, **c**, H5-SS-, and **d**, H1-SS-specific IgG titers over time. Endpoint (week 12) IgG titers are plotted on the right for comparison. **e**, nsEMPEM analyses for pooled antisera raised against H5 HA_Phe_ or reoH5HA_Phe_. 3D composite maps and representative 2D class averages generated from negative-stain reconstructions of polyclonal immune complexes with H1 HA (CA/09) are shown for each group, with dotted lines highlighting the likely position of the Fab population at low abundance. **f**, Body weight loss and **g**, Kaplan-Meier survival curves of mice after lethal challenge (10× mLD_50_) with the 2009 H1N1 Pandemic virus (H1N1pdm09, A/Netherlands/602/2009). **h**, A prime-boost immunization *via* intramuscular injection in female BALB/c mice. All mice were primed with FZ (20 µL per mouse) adjuvanted with CpG (10 µg per mouse) and then boosted with PBS (*n =* 12), FZ adjuvanted with CpG (*n =* 16), H5 HA_Phe_ (1 µg per mouse) adjuvanted with alum/CpG (100/25 µg per mouse, *n =* 15) twice, or reoH5HA_Phe_ (1 µg per mouse) adjuvanted with alum/CpG (100/25 µg per mouse, *n =* 16) twice. **i**, H1-SS-specific IgG titers over time. **j**, Body weight loss and **k**, Kaplan-Meier survival curves of mice after lethal challenge (10× mLD_50_) with the H1N1 virus (PR8/34). Arrows indicate prime and boost immunizations in **b-d**. Data are presented as geometric mean ± s.d. of the log_10_-transformed values in **b-d**,**i** or mean ± s.d. in **f**,**j**. Dashed lines indicate the limit of quantification in **b-d**,**i** or 25% weight loss in **f**,**j**. Comparison of two groups was performed using the two-tailed Mann-Whitney U-test in **b-d**. Comparison of multiple groups was performed using one-way ANOVA followed by a Tukey’s post-hoc test in **i**. Comparison of survival curves was performed using a pairwise log-rank test in **g,k**. P values of 0.05 or less were plotted and considered significant.

We also evaluated reoH5HA_Phe_ as a booster immunization in FZ-vaccinated mice. Since a single booster with H5 HA_Phe_ or reoH5HA_Phe_ likely induced a naïve rather than anamnestic response in FZ-pre-immune mice, we performed two boosts with H5 HA_Phe_ or reoH5HA_Phe_ followed by lethal challenge with two H1N1 viruses, H1N1pdm09 and A/Puerto Rico/8/1934 (PR8/34) (**Fig. 5h**). Similar to previous results (**Fig. S6e**), boosting with FZ elevated IgG titers to seasonal influenza HAs (**Fig. S8a**) with limited stem-directed responses (**Fig. 5i, Fig. S8b**) and minimal cross-reactivity to H5 HA (**Fig. S8c**). Compared with H5 HA_Phe_, boosting with reoH5HA_Phe_ substantially enhanced H5 HA-specific neutralizing antibody responses (**Fig. S8d**) and balanced, stem-directed IgG titers (**Fig. S8e**) with increased cross-reactivity to H1 HAs (**Fig. S8f,g**). When challenged with H1N1pdm09, a single dose of FZ provided limited protection (1/6 mice), whereas booster immunizations with FZ, H5 HA_Phe_, or reoH5HA_Phe_ all mitigated weight loss and provided complete protection (**Fig. S8h,i**), despite negligible cross-neutralizing activity against the H1N1 virus (**Fig. S8j**). When challenged with PR8/34, boosting with H5 HA_Phe_ improved survival (4/8 mice) relative to the PBS control (2/6 mice), while reoH5HA_Phe_ protected 7/8 mice almost equivalent to FZ (**Fig. 5j,k**) in the absence of cross-neutralizing activity against PR8/34 (**Fig. S8k)**. Collectively, these results show that reoH5HA_Phe_ enhanced stem-directed antibody responses against H1 HAs and conferred heterosubtypic protection against H1N1 viruses.

## Discussion

Human infections with HPAI H5N1 viruses have historically resulted in high mortality, and the continuous circulation and viral evolution in animal reservoirs may generate variants distinct from available vaccine strains, against which human populations lack pre-existing immunity(1–3, 7). Although many vaccine candidates have been developed(8–15), including mRNA vaccines for rapid pandemic responses, they primarily elicit neutralizing antibodies to the immunodominant and variable HA-head with decreased cross-reactivity to mismatched strains. Two to three immunizations with the licensed H5N1 vaccine (based on VN/04) were needed to induce cross-reactive antibodies to clade 2.3.4.4b H5 HA in humans(13). mRNA vaccines based on clade 2.3.4.4b H5 HA(9, 15, 16) or stabilization of this HA(12) enhanced neutralizing antibody responses and protected against homologous lethal challenge, although protection against heterologous strains remains suboptimal or unclear. In addition, antigenic mapping enabled identification of antigenically central H5 HAs to confer heterologous immunity following prime-boost immunizations(10). Considering the difficulty of predicting genetic sequences of emerging viruses, effective pandemic preparedness necessitates vaccine candidates that establish broadly protective immunity on conserved HA epitopes(5).

In contrast to HA-head, HA-stem is evolutionarily conserved and targeted by bnAbs(22–26), providing structural and functional basis for stem-based vaccine strategies. In this study, we employed the antigen-reorientation approach to overcome HA immunodominance and guide antibody responses toward stem epitopes. We designed reoH5HA_Phe_ based on the historical VN/04 sequence and showed that reoH5HA_Phe_ adjuvanted with the optimized alum/CpG adjuvant enhanced robust stem-directed responses with balanced IgG1 and IgG2 subclasses. In naïve mice, a single immunization with reoH5HA_Phe_ conferred 50% or more protection against heterologous H5N1 (TX/24) lethal challenge, whereas H5 HA_Phe_ failed to protect. nsEMPEM analysis showed that, while both H5 HA_Phe_ and reoH5HA_Phe_ induced cross-reactive antibodies against the VE epitope, only reoH5HA_Phe_ elicited those targeting two stem epitopes on the heterologous H5 HA (TX/24), contributing to improved protection. In addition, in FZ-pre-immune mice, a single booster immunization with reoH5HA_Phe_ conferred complete heterologous protection with reduced body weight loss and viral replication in respiratory tissues, while boosting with FZ or H5 HA_Phe_ provided minimal to no protection. In both studies, reoriented antigen presentation and alum binding collectively enhanced the stem immunogenicity of reoH5HA_Phe_. While cross-reactive antibodies against the VE and anchor epitopes were more abundant than those against the central stem epitope in the single-dose immunization, we only observed antibody binding to the central stem epitope in FZ-pre-immune mice boosted with reoH5HA_Phe_. Since nsEMPEM was performed using antisera two weeks post-boost, it remained to be determined whether cross-reactive antibodies targeting VE or anchor epitopes were elicited at later times and whether reoH5HA_Phe_ stimulated a naïve or recall response toward the central stem epitope. Despite limited cross-neutralizing activity against TX/24, increased stem-directed responses with balanced IgG1 and IgG2 subclasses correlated with reduced viral titers and improved survival, suggesting protection *via* Fc-mediated mechanisms, similar to previous studies with stem-only immunogens(38). Since HA-stem is considered mutationally intolerant(47), the cross-clade protection suggests that reoH5HA_Phe_ has potential to complement the antigenic matching strategy and serve as a single-dose vaccine candidate to protect against antigenically mismatched H5N1 viruses in a pandemic setting.

Besides HA, neuraminidase (NA) is another influenza viral glycoprotein and target for neutralizing antibodies(48). Although natural infection often induces robust NA-specific immunity, seasonal vaccines rarely elicit NA-reactive B cells or antibodies due to poor display of critical NA epitopes(49). Insufficient NA-specific immunity from the prime-boost immunization with FZ may explain why it failed to protect against the H5N1 challenge, despite the presence of N1 NA in both FZ and the challenge virus. When the immunogenicity of NA in the split virus vaccine was retained through optimized bioprocessing(50), cross-reactive NA antibodies were critical for heterologous protection(51). Therefore, enhancing responses against conserved epitopes on both HA-stem and NA may further expand the protective breadth against antigenically distinct influenza viruses(5, 52, 53).

In addition to H5 subtypes, we showed that immunizations with reoH5HA_Phe_ induced significantly higher antibody responses against H1-SS than H5 HA_Phe_ and protected against heterosubtypic H1N1 viruses in both naïve and pre-immune mice. While H5 HA-specific IgG titers were comparable between the two groups, reoH5HA_Phe_ significantly enhanced stem-directed responses that correlated with improved survival, in the absence of cross-neutralizing activity. Since protein subunit vaccines generally induce poor T cell responses(54), cross-reactive antibodies elicited by reoH5HA_Phe_ most likely mediated protection against H1N1 viruses, in contrast to previous studies using viral vectors expressing H5 HA(55, 56), which induced robust CD8^+^ T cell responses to achieve heterosubtypic H1N1 protection. Future passive transfer studies will allow examination of this potential immune correlate of protection. Whereas antibodies raised against H5 HA_Phe_ disrupted HA trimers and bound to internal epitopes in the nsEMPEM analysis, they provided minimal protection against viral challenge. By contrast, antibodies from the reoH5HA_Phe_ group recognized distinct epitopes on the intact H1 HA trimer and protected against the H1N1 challenge, suggesting that reoH5HA_Phe_ binding to alum stabilized the trimer conformation, reduced exposure of internal epitopes, and enhanced the quality of antibody responses. In addition, at least after a single immunization, reoH5HA_Phe_ elicited a higher fraction of anchor-than central stem-directed antibodies from the nsEMPEM analysis, potentially resulting in lower H1-SS-specific IgG titers relative to H5-SS. Further increasing H1-SS-cross-reactive responses may mitigate the initial weight loss and achieve complete protection against H1N1 challenge. This difference in stem-directed titers also suggests that although HAs from different subtypes share similar stem antigenicity, their stem immunogenicity may differ, warranting further investigations to define epitope specificity elicited by different HA stems.

We also acknowledge limitations of our studies. First, the current antigen reorientation strategy uses the Th2-biased alum adjuvant and is not compatible with the mRNA vaccine platform, although combining with Th1-biased adjuvants (*e.g.*, CpG) could induce a balanced Th1/Th2 response(57). Second, while the current study was conducted in mice for proof-of-concept, the protective efficacy and breadth remain to be tested in other animal models. A single immunization with the replicating RNA vaccine encoding H5 HA (VN/04) protected 2/6 mice against bovine H5N1 challenge but fully protected nonhuman primates against the challenge following a prime-boost regimen(14), suggesting that the protective efficacy varies across animal models. Third, previous studies found that H5N1 vaccination induced antibody and memory B cell responses to HA-stem in humans(58), whereas H5 HA_Phe_ or reoH5HA_Phe_ likely induced a naïve response in pre-immune mice, potentially because of our simplified pre-existing immunity model and differences between mouse and human repertoires. It remains to be determined whether reoH5HA_Phe_ induces a naïve or recall response toward HA-stem in human organoid systems(59). Finally, while we focused on characterizing antibody responses, detailed analysis of antibody-dependent cellular cytotoxicity(60, 61), complement-mediated enhancement(62), and cellular immunity(63) would collectively define immune correlates of protection.

In summary, we designed reoH5HA_Phe_ as an H5 pandemic influenza vaccine candidate that immunofocuses antibody responses on HA-stem and protects against H5N1 and H1N1 viruses in both naïve and pre-immune animals. The antigen reorientation approach also has potential to accelerate development of universal vaccines and establish broad immunity against antigenically distinct, emerging viruses.

## Materials and Methods

### Cell lines

Expi293F cells (Thermo Fisher) were maintained in Freestyle293/Expi293 medium (2:1, *v/v*, Thermo Fisher) in polycarbonate shaking flasks (Triforest Labware). MDCK-SIAT1 cells (Cytion) were maintained in DMEM (Cytiva) supplemented with 10% FBS (GeminiBio) and 1% penicillin-streptomycin-L-glutamine (Corning). MDCK-SIAT1 culture medium was supplemented with 1 mg mL^-1^ G418 (Thermo Fisher). The MDCK cell line used for plaque assay of authentic viruses was adapted from its original source in our laboratory. Cells were grown at 37 °C and 5% CO_2_ in Eagle’s minimum essential medium (MEM) containing 5% newborn calf serum and were monitored routinely for mycoplasma contamination. Stellar competent cells were purchased from Takara Bio.

### Antibodies

HA-specific mAbs (H5.3, H5M9, MEDI8852, CR9114, FI6v3, 222-1C06, and CH65) were expressed in Expi293F cells *via* transient transfection. Goat anti-mouse IgG, HRP-conjugated (clone Poly4053, BioLegend #405306), goat anti-mouse IgG1-HRP (SouthernBiotech #1070-05), goat anti-mouse IgG2a-HRP (SouthernBiotech #1083-05), goat anti-mouse IgG2b-HRP (SouthernBiotech #1093-05), goat anti-rabbit IgG, HRP (SouthernBiotech #4030-05), rabbit anti-human IgG, HRP-conjugated (Abcam #ab6759), goat anti-human IgG-Fc, HRP-conjugated (SouthernBiotech #2048-05 and #2014-01), streptavidin-HRP (Pierce #21130) were used as detecting antibodies for Western blot analysis or enzyme-linked immunosorbent assays (ELISAs). Rabbit anti-influenza A nucleoprotein antibody (clone HL1078, Invitrogen, MA5-42363) was used for influenza A virus microneutralization assays.

### Antigen and antibody cloning

DNA encoding H5 HA (A/Vietnam/1203/2004 or VN/04) was cloned into the pADD2 vector with a foldon trimerization domain followed by a 6×His tag on the C-terminus. OligoD (12 units) was inserted into the 150-loop of H5 HA after residue N164 (based on H3 numbering) to create reoH5HA. Phe45_HA2_ substitution was introduced to both H5 HA and reoH5HA. DNAs encoding HA ectodomains of A/New Caledonia/20/99 (NC/99), A/California/07/2009 (CA/09), A/Japan/305/1957 (JP/57), VN/04, A/Victoria/4897/2022 IVR-238 (VIC/22), A/Darwin/9/2021 SAN-010 (DRW/21), B/Phuket/3073/2013 (B/HA PKT/13), B/Michigan/01/2021 (B/HA MI/21), A/Cygnus columbianus/Hubei/50/2020 (HB/20), A/duck/Bangladesh/58330/2023 (BD/23), A/American Wigeon/South Carolina/22-000345-001/2021 (SC/21), A/Texas/37/2024 (TX/24), H1-SS, H5-SS, and Ebola glycoprotein (GP) with the mucin-like domain deleted (residues 1-308 and 491-656) were cloned into the pADD2 vector with a GCN4 trimerization domain followed by an Avi-Tag and a 6×His tag on the C-terminus. DNA encoding HA ectodomain of TX/24 fused to *H. pylori* ferritin, mFcγRIIIa fused to a GCN4 dimerization domain and an Avi-Tag, and mFcγRIV fused to VRC01-Fc were cloned into the pADD2 vector. DNA fragments encoding the variable heavy chain (HC) and light chain (LC) of antibodies were inserted into an expression plasmid encoding VRC01 HC and LC constant domains. All plasmid sequences were confirmed by Sanger (Functional Biosciences) or whole-plasmid sequencing (Plasmidsaurus). For transfection, plasmids were transformed into Stellar cells, isolated by Maxiprep kits (NucleoBond Xtra Maxi kit, Macherey Nagel), filtered through a sterile 0.45-µm membrane in a biosafety cabinet and stored at -20 °C.

### Protein expression and purification

All antigens and antibodies were expressed in Expi293F cells. Expi293F cells were cultured at 37 °C under constant shaking (120 rpm) in a humidified CO_2_ (8%) incubator. Expi293F cells were transfected at a density of 3-4 × 10^6^ cells mL^-1^. For 200 mL transfection, the transfection mixture was made by adding 160 µg plasmid DNA to 20 mL expression medium, followed by dropwise addition of 260 µL FectoPro transfection reagent (Polyplus) with vigorous mixing. For antibody production in 200 mL Expi293F cells, the transfection mixture contained 100 µg LC plasmid DNA and 100 µg HC plasmid DNA. Transfection mixtures were incubated at room temperature for 10 min before being transferred to Expi293F cells. *D*-glucose (4 g/L, Sigma-Aldrich) was added immediately post-transfection. Cells were boosted with valproic acid (3 mM, Acros Organics) 24 hours post-transfection and harvested on day 4 or 5 by centrifugation at 7000 ×*g* for 5 min. The supernatant was filtered through a 0.45-µm membrane for subsequent purification. Biotinylated proteins were expressed using the same protocol but in the presence of the BirA enzyme.

Antigen proteins with 6×His tags were purified with HisPur Ni-NTA resin (Thermo Fisher). Filtered supernatant from Expi293F cells was mixed with Ni-NTA resin (10 mL resin per liter supernatant) and incubated at 4 °C overnight. The mixture was passed through a gravity-flow column, washed with 20 mM imidazole in HEPES buffer (HBS, 20 mM HEPES, pH 7.4, 150 mM NaCl), and eluted with 300 mM imidazole in HBS. Elution was concentrated with centrifugal filters (30 or 50 kDa MWCO, Millipore Sigma) for size-exclusion chromatography using a Superose 6 (Increase 10/300 GL, Cytiva) or Superdex 200 column (Increase 10/300 GL, Cytiva) on an ÄKTA Pure Protein Purification System (Cytiva). Peak fractions were pooled, concentrated, buffer-exchanged to HBS with 10% glycerol and filtered through a 0.22-µm membrane.

All antibodies and proteins with Fc were purified with Protein A chromatography. Filtered supernatant from Expi293F cells was directly applied to a MabSelect SuRe column (Cytiva) on an ÄKTA System. The column was washed with HBS and then antibodies were eluted with glycine (100 mM, pH 2.8) to HEPES buffer (1 M, pH 8.0). Fractions were concentrated and buffer-exchanged to HBS with 10% glycerol.

Protein concentration was determined by absorbance at 280 nm (A_280_), and purity was assessed by protein gel electrophoresis. Protein samples were flash-frozen in liquid nitrogen and stored at -80°C.

### Size-exclusion chromatography-multi-angle light scattering (SEC-MALS) analysis

SEC-MALS analysis was performed on a 1260 Infinity II high-performance liquid chromatography system (Agilent) coupled with a miniDAWN and Optilab detectors (Wyatt Technologies) for light scattering and refractive index analysis. Purified proteins (0.2 to 0.5 mg mL^-1^) were loaded onto a Superdex 200 column (Increase 3.2/300, Cytiva) sequentially for analysis. ASTRA software (version 7.3.2.21, Wyatt Technologies) was used for quantitative analysis of the M_w_ of antigen proteins.

### Differential scanning fluorimetry

Thermal melting profiles of proteins were measured by differential scanning fluorimetry on a CFX Connect Real-Time PCR Detection System (Bio-Rad). Protein samples (0.1 mg mL^-1^) were mixed with SYPRO Protein Gel Stain (5×, Thermo Fisher) and transferred to a 96-well qPCR plate (USA Scientific). Samples were then subjected to a temperature gradient from 22 to 95 °C at a heating rate of 1 °C per 10 seconds. Thermal melting curves were plotted using the normalized first derivative of the relative fluorescence unit (RFU). Melting temperatures were defined as peaks in the thermal melting curves.

### Biolayer interferometry (BLI)

BLI experiments were performed on an OctetRed 96 system (FortéBio). All samples were diluted with the Octet buffer (PBS with 0.02% Tween-20 and 0.1% bovine serum albumin), and assays were performed under agitation (1000 rpm). Biotinylated antigens (*e.g.*, H5 HA, reoH5HA, H5 HA_Phe_, and reoH5HA_Phe_ at 500 nM) were loaded onto Streptavidin biosensors (Sartorius) and then dipped into antibody solutions (*e.g.*, MEDI8852, CR9114, FI6v3, H5M9, or CH65 at 200 nM) for binding analysis followed by dissociation in the Octet buffer. Data were processed by Data Analysis software (version 9.0.0.15, FortéBio) and then plotted.

### Antigen-alum binding assays

H5 HA, reoH5HA, H5 HA_Phe_, and reoH5HA_Phe_ were first incubated with alum (Alhydrogel, InvivoGen, protein/alum, 1/10 to 1/100, *w/w*) for 30 min in HBS at room temperature and then CpG adjuvant (ODN 1018, MedChemExpress) was added to the mixture at various ratios (CpG/protein, 1/1 to 1/25, *w/w*). After 30-min incubation, naïve mouse serum was added to the mixture to a final concentration of 10% (*v/v*). The mixture with alum alone served as the control. Mixtures were further incubated at 37 °C under constant shaking (220 rpm) on an orbital shaker for 24 hours before being centrifuged to pellet alum (10,000 ×*g* for 5 min). Alum pellets were rinsed with PBS and re-pelleted (three times) by centrifugation, then resuspended in SDS-PAGE sample loading buffer for Western blot analysis to determine the amount of alum-bound protein. Resuspended alum pellets were boiled at 95 °C for 10 min before being applied to a polyacrylamide gel (4-20% Mini-PROTEAN, Bio-Rad). Proteins were then transferred to a polyvinylidene difluoride membrane using the Trans-Blot Turbo transfer system (Bio-Rad). Blots were blocked in PBST with 10% non-fat dry milk (Bio-Rad), incubated with rabbit anti-His-tag polyclonal antibody (1:4000 in PBST with 10% non-fat dry milk) and then detected with goat anti-rabbit IgG, HRP-conjugated (1:4000 in PBST with 10% non-fat dry milk). Luminescent signals were developed using a luminol-based substrate (Pierce ECL, Thermo Fisher) for imaging on a chemiluminescence imager (ChemiDoc Touch System, Bio-Rad).

The supernatant was collected for either gel electrophoresis followed by Western blotting or ELISA to measure the concentration of unbound protein antigens. For the ELISA measurement, Nunc 96-well MaxiSorp plates (Thermo Fisher) were coated with the antigen-binding fragment (Fab) of MEDI8852 (4 µg mL^-1^ in PBS, 60 µL per well) for 1 hour at room temperature. Plates were washed 3 times with Milli-Q H_2_O (300 µL) using a plate washer (ELx405 BioTek) and then blocked with ChonBlock (120 µL per well, Chondrex) overnight at 4 °C. For subsequent steps, all dilutions were made in the Octet buffer, and ELISA plates were rinsed with PBST (300 µL, 3 times) in between steps. Antigens at appropriate dilutions were added to the plate and then detected by H5M9 IgG (1:4000). Rabbit anti-human IgG-Fc, HRP-conjugated (1:4000) was added for 1-hour incubation before rinsing with PBST 6 times. ELISA plates were developed with the TMB (3,3’,5,5’-tetramethylbenzidine) substrate (1-Step Turbo-TMB, Thermo Fisher) for 6 min and terminated with sulfuric acid (2 M). Absorbance at 450 nm was recorded on a microplate reader (Synergy HT, BioTek). reoH5HA_Phe_ with a series of known concentrations (5-fold dilution starting from 2 µg mL^-1^) was used to establish a standard curve, and the amount of antigen was quantified by fitting the absorbance values to this curve.

### Streptavidin- and alum-based ELISAs

For streptavidin-based ELISAs, Nunc 96-well MaxiSorp plates were coated with streptavidin (4 µg mL^-1^ in PBS, 60 µL per well) for 1 hour at room temperature. Plates were washed 3 times with Milli-Q H_2_O and then blocked with ChonBlock (120 µL per well) overnight at 4 °C. For subsequent steps, all dilutions were made in the Octet buffer, and ELISA plates were rinsed with PBST in between steps. C-terminally biotinylated H5 HA, reoH5HA, H5 HA_Phe_, and reoH5HA_Phe_ (2 µg mL^-1^) were added to the plates and incubated for 1 hour at room temperature. mAbs (H5.3, H5M9, MEDI8852, and FI6v3) were serially diluted (10-fold dilution starting from 20 nM) and then added to the ELISA plates for 1-hour incubation at room temperature. Rabbit anti-human IgG, HRP-conjugated (1:4,000) was added for 1-hour incubation before rinsing with PBST 6 times. ELISA plates were developed with the TMB substrate for 5 min and terminated with sulfuric acid. Absorbance at 450 nm was recorded on a microplate reader.

For alum-based ELISAs, Nunc 96-well MaxiSorp plates were coated with Ebola GP with the C-terminal oligoD insertion (4 µg mL^-1^ in PBS, 60 µL per well) for 1 hour at room temperature. These plates were washed 3 times with Milli-Q H_2_O and then blocked with ChonBlock overnight at 4 °C. Alum (100 µg mL^-1^ in HBS) was added to the plates and incubated for 1 hour at room temperature. After rinsing, H5 HA, reoH5HA, H5 HA_Phe_, and reoH5HA_Phe_ (2 µg mL^-1^) were added to the plates and incubated for 1 hour at room temperature. Similar to streptavidin-based ELISAs, serially diluted mAbs were added to the plates and detected by rabbit anti-human IgG, HRP-conjugated (1:4,000).

### Mouse immunization studies

All animals were maintained in accordance with the Public Health Service Policy (PHS) for Humane Care and Use of Laboratory Animals under a protocol (#A006864) approved by the University of Wisconsin-Madison Institutional Animal Care and Use Committee (IACUC). Female BALB/c mice (6-8 weeks) were purchased from Jackson Laboratories, maintained with 12-hour light/dark cycles at 65-75 °F with 40-60% humidity, and randomly grouped for the studies. Antigens were mixed with adjuvants before injections, with the total volume adjusted to 100 µL with HBS. Injection mixtures were incubated at room temperature for 30 min before use. All mice were immunized *via* intramuscular injections. To compare the immunogenicity of H5 HA_Phe_ and reoH5HA_Phe_, we immunized two groups of BALB/c mice (n = 10) with 5 µg antigens adjuvanted with 150/5 µg alum/CpG. To screen different adjuvant combinations, we immunized 6 groups of BALB/c mice (n = 5) with 5 µg reoH5HA_Phe_ adjuvanted with 100 µg alum and 10 µg of an alum-compatible adjuvant, including CpG, monophosphoryl lipid A or MPLA, 3M-052, polyinosinic:polycytidylic acid or poly(I:C), and globotriaosylceramide or Gb3. To compare the protective efficacy of H5 HA_Phe_ and reoH5HA_Phe_, we performed a single-dose immunization in 4 groups of BALB/c mice (n = 8) with 5 µg antigens adjuvanted with 100/10 µg alum/CpG or 1 µg antigens adjuvanted with 100/25 µg alum/CpG. We also performed a prime-boost immunization in another two groups of BALB/c mice (n = 16) with 5 µg H5 HA_Phe_ or reoH5HA_Phe_ adjuvanted with 100/10 µg alum/CpG. To simulate pre-existing immunity, we primed 4 groups of BALB/c mice with FZ (20 µL, Quadrivalent, 2023-2024 Formula, BEI Resources) adjuvanted with CpG (10 µg) and then boosted them with PBS (n = 6), FZ adjuvanted with CpG (10 µg) (n = 10), H5 HA_Phe_ (1 µg) adjuvanted with alum/CpG (100/25 µg) (n = 10), or reoH5HA_Phe_ (1 µg) adjuvanted with alum/CpG (100/25 µg) (n = 10). Single-dose and FZ-priming immunization studies were repeated to assess viral replication post-lethal challenge (n = 6 per group). To evaluate the heterosubtypic protection against H1N1 viruses, we performed a three-dose immunization in two groups of BALB/c mice (n = 16) with 1 µg H5 HA_Phe_ or reoH5HA_Phe_ adjuvanted with 100/25 µg alum/CpG. Mice were immunized with PBS (n = 10) or H1 HA (CA/09) (n = 12) as controls. To simulate pre-existing immunity, we primed 4 groups of BALB/c mice with FZ (20 µL) adjuvanted with CpG (10 µg) and then boosted them with PBS (n = 12), FZ adjuvanted with CpG (10 µg) (n = 8), H5 HA_Phe_ (1 µg) adjuvanted with alum/CpG (100/25 µg) twice (n = 15), or reoH5HA_Phe_ (1 µg) adjuvanted with alum/CpG (100/25 µg) twice (n = 16).

Pre-immune, interim and final blood samples were collected by retro-orbital or submental bleeding into serum gel tubes (Sarstedt). Serum gel tubes were centrifuged at 10,000 ×*g* for 6 min, and sera were collected and stored at -80 °C.

### Serum ELISAs

We used proteins with a different trimerization domain from the immunogens for ELISA binding analysis of antisera. To deplete serum antibodies against C-terminal tags (*e.g.*, 6×His tags), we added GFP with a 6×His tag (2 µg mL^-1^, final concentration) to the ELISA dilution buffer. Nunc 96-well MaxiSorp plates were coated with streptavidin for 1 hour at room temperature. These plates were washed 3 times with Milli-Q H_2_O using a plate washer and then blocked with ChonBlock overnight at 4 °C. Biotinylated antigens (2 µg mL^-1^) were added to the plates and incubated for 1 hour at room temperature. Mouse antisera were serially diluted (5-or 10-fold dilution) and then added to the ELISA plates for 1-hour incubation at room temperature. Goat anti-mouse IgG, HRP-conjugated (1:4000) was added for 1-hour incubation before rinsing with PBST 6 times. ELISA plates were developed with the TMB substrate for 6 min and terminated with sulfuric acid. Absorbance at 450 nm was recorded on a microplate reader. To test antigen-specific antibody responses, ELISA plates were incubated with biotinylated HAs from the following strains: VN/04, H1 VIC/22, H3 DRW/21, B/HA PKT/13, B/HA MI/21. To test stem-specific antibody responses, ELISA plates were incubated with biotinylated H1-SS or H5-SS. To test the cross-reactivity to different HAs, ELISA plates were incubated with biotinylated HAs from the following strains: NC/99, H1 CA/09, JP/57, HB/20, BD/23, SC/21, or TX/24. For competition ELISA assays, competing antibodies (*e.g.*, MEDI8852 or 222-1C06, 100 nM) were preincubated on the plate with antigens for 1 hour before adding serially diluted mouse antisera.

### Mouse Fcγ receptor (mFcγR) binding assays

Antigen-specific IgG binding to mFcγRIIIa or mFcγRIV was used as a correlate of effector function activation. Nunc 96-well MaxiSorp plates were coated with H5 HA (TX/24) displayed on *H. pylori* ferritin nanoparticles (4 µg mL^-1^) for 1 hour at room temperature. Plates were washed 3 times with Milli-Q H_2_O using a plate washer and then blocked with ChonBlock overnight at 4 °C. Mouse antisera were serially diluted (3-fold dilution) and then added to the ELISA plates for 1-hour incubation at room temperature. mFcγRIIIa-GCN4-Avi (biotinylated) or mFcγRIV-VRC01-Fc (5 µg mL-1) was added for 1-hour incubation before detection with streptavidin-HRP (1:10,000) or goat anti-human IgG Fc, HRP-conjugated (1:4000), respectively. ELISA plates were rinsed with PBST 6 times, developed with the TMB substrate and terminated with sulfuric acid. Absorbance at 450 nm was recorded on a microplate reader.

### Purification and digestion of IgG from pooled antisera

Pooled antisera from each group were filtered with 0.22-µm membrane and directly applied to a HiTrap Protein G HP column (Cytiva) on an ÄKTA System. The column was washed with HBS and then polyclonal IgG was eluted with glycine (100 mM, pH 2.2) to HEPES buffer (1 M, pH 8.0). Fractions were concentrated and buffer-exchanged to HBS with 10% glycerol. IgG concentration was determined by A_280_, and its purity was assessed by protein gel electrophoresis. IgG was digested with a mixture of immobilized ficin (5 µL per mg IgG, Thermo Fisher) and papain (1 µL per mg IgG, Thermo Fisher) for 2 hours at 37 °C in the presence of cysteine (25 mM) and 2-mercaptoethylamine (50 mM). Digestion was quenched with iodoacetamide (100 mM) in Tris buffer (100 mM, pH 7.5) and products were purified with Zeba desalting columns (Thermo Scientific). Purity of the digestion mixture was assessed by size-exclusion chromatography on a Superdex 200 column (Increase 10/300 GL, Cytiva).

### Negative-stain electron microscopy polyclonal epitope mapping (nsEMPEM)

HA (10 µg) was complexed with polyclonal Fab at 1/20 to 1/40 ratios (*w/w*) overnight at 4 °C before purification on a Superose 6 column (Increase 3.2/300 GL, Cytiva). Fractions containing HA-Fab immune complexes were pooled and deposited for 30 seconds at approximately 8-12 µg mL^-1^ on freshly glow-discharged, carbon-coated copper grids (400 mesh, Electron Microscopy Sciences). Grids were then washed 3 times with Milli-Q H_2_O followed by staining with uranyl formate (2%, *w/v*). Images were acquired at 73,000× magnification (2.05 Å/pixel) on a Talos F200C microscope (Thermo Fisher) operating at 200 kV, equipped with a BM-Ceta camera. Electron micrographs were collected in an automated fashion using SerialEM (Thermo Fisher) before processing in cryoSPARC (v5.0.6) as previously described. Composite figures were created by docking 20-Å low-pass filtered Fab volumes into 3D density corresponding to bound Fab for each complex, while HA is represented as a 20-Å low-pass filtered volume. When evidence for a Fab was only present in 2D class averages, the presumed position, based on similar 2D class averages from known HA-Fab complexes, in composite figures was marked by dotted lines around the Fab. UCSF ChimeraX (version 1.10) was used to generate composite figures.

### Production and titration of H5N1 pseudoviruses

Pseudotyped lentiviruses encoding a luciferase-ZsGreen reporter were produced in Expi293F cells by co-transfection of 6 plasmids, including a packaging vector (pHAGE-Luc2-IRES-ZsGreen), a plasmid encoding full-length H5 HA (VN/04 or TX/24), a plasmid encoding full-length neuraminidase (NA, A/American green-winged teal/Washington/2014), and 3 helper plasmids (pHDM-Hgpm2, pHDM-Tat1b, and pRC-CMV_Rev1b). Expi293F cells were adjusted to 3 × 10^6^ cells per mL before transfection. For 10 mL of Expi293F cells, transfection mixture was prepared by adding 6 plasmids (5 µg packaging vector, 1.25 µg plasmid encoding H5 HA, 0.25 µg plasmid encoding NA, 1.25 µg of each helper plasmid) to 1 mL culture medium, followed by the addition of 15 µL BioT transfection reagent (Bioland Scientific) in a dropwise manner with vigorous mixing. After 10-min incubation at room temperature, the transfection mixture was transferred to Expi293F cells and cells were immediately boosted with *D*-glucose (4 g/L). One day post-transfection, cells were pelleted at 300 ×*g*, resuspended in fresh culture medium and boosted with *D*-glucose (4 g/L) and valproic acid (3 mM). Pseudoviruses were harvested 3 days post-transfection, filtered through a 0.45-µm membrane, and incubated with TPCK-treated trypsin (2 µg mL^-1^, Thermo Fisher) at 37 °C for 45 min on an orbital shaker (120 rpm). Trypsinization was quenched by adding FBS to the mixture (5% of total volume). After processing, pseudoviruses were aliquoted, flash-frozen in liquid nitrogen, and stored at -80 °C. For titration, serially diluted pseudoviruses were used to infect MDCK-SIAT1 cells. Oseltamivir (1 µM), amphotericin B (2.5 µg mL^-1^), and polybrene (10 µg mL^-1^) were supplemented to the pseudovirus to increase infectivity. After 48-hour incubation, medium was removed and 70 µL of luciferase substrates (BriteLite Plus, Perkin Elmer) were added to each well. Luminescent signals were recorded on a microplate reader (Tecan M200). For neutralization assays, pseudoviruses were diluted in culture medium to yield more than 10^6^ relative luminescence units (RLU).

### H5N1 pseudoviral neutralization assays

Antisera were first treated with receptor-destroying enzymes (Denka Seiken) at 37 °C for 18 h before adding sodium citrate (2.5%, *w/v*) to quench the treatment (antisera/receptor-destroying enzymes/sodium citrate = 1/3/3, *v/v*). Antisera were then heat inactivated at 56 °C for 60 min and passed through centrifugal filters (Corning, 0.22 µm). Serially diluted antisera were incubated with H5N1 pseudoviruses (supplemented with 1 µM Oseltamivir, 2.5 µg mL^-1^ amphotericin B, and 10 µg mL^-1^ polybrene) for 1 hour before being transferred to MDCK-SIAT1 cells for further incubation. After 48 hours, medium was removed and 70 µL of luciferase substrate was added to each well. Luminescent signals were recorded on a microplate reader (Tecan M200). Percent infection was normalized to cells only (0% infection) and virus only (100% infection) on each plate. Neutralization titers (NT_50_) were calculated as the serum dilution where a 50% inhibition of infection was observed. Neutralization assays were performed in technical duplicates.

### Influenza A microneutralization assays

Viral infectivity was titrated in MDCK-SIAT1 cells, and 100× 50% tissue culture infectious dose (TCID_50_) was used for influenza A virus microneutralization assays. Antisera were processed with receptor-destroying enzymes and sodium citrate, heat inactivated and passed through 0.22-µm centrifugal filters before use. Serially diluted antisera were incubated with wild-type viruses (2 µg mL^-1^ TPCK-trypsin-treated) for 1 hour before being transferred to MDCK-SIAT1 cells. After 48 hours, medium was removed from all wells, rinsed cells with PBS twice and fixed cells with paraformaldehyde (BioLegend). After fixation, cells were rinsed with PBS twice, permeabilized with Triton X-100 (Cell Signaling Technology), and stained with rabbit anti-influenza A nucleoprotein antibody (1:4,000) for 1 h, followed by goat anti-rabbit IgG, HRP-conjugated (1:4,000, 1 h). Plates were rinsed with PBST 6 times and then developed with the TMB substrate and terminated with sulfuric acid. Absorbance at 450 nm was recorded on a microplate reader. Percent infection was normalized to cells only (0% infection) and virus only (100% infection) on each plate. Microneutralization assays were performed in technical duplicates.

### Lethal challenge studies

Experiments with highly pathogenic avian influenza (HPAI) H5N1 virus were conducted in Biosafety Level 3 (BSL-3) containment laboratories at the Influenza Research Institute, University of Wisconsin-Madison, which is approved by the Federal Select Agent Program for studies with these viruses. All experiments were approved by the Institutional Biosafety Committee of the University of Wisconsin-Madison, and all animal experiments were approved by the University of Wisconsin-Madison School of Veterinary Medicine Institutional Animal Care and Use Committee (protocol #V006426). Immunized BALB/c mice were anesthetized by isoflurane and infected by intranasal inoculation (25 µL total volume, equally between each nostril) with the indicated virus at 10 or 15× mLD_50_ diluted in PBS. Mice were monitored at least daily for signs of illness, and body weights were recorded for up to 2 weeks. Mice were euthanized if they lost 25% or more of their initial body weight or could not remain upright. To assess viral replication, mice were euthanized 3 days post-infection for extraction of lung, nasal turbinate, and brain tissues. Tissues were homogenized, clarified, and then applied to confluent MDCK cell monolayers overlaid with 1× MEM containing 0.3% bovine serum albumin, 2% low melting point agarose, and 0.6 µg mL^-1^ L-(tosylamido-2-phenyl) ethyl chloromethyl ketone (TPCK)-treated trypsin to determine viral titers.

### Statistics and reproducibility

Western blot analysis and serological analyses were repeated at least twice with similar results, and one representative data set is presented. Statistical analyses were performed using GraphPad Prism software (version 10.6.1). Non-transformed data are presented as mean ± SD. ELISA titers, NT_50_, and viral titers were log_10_-transformed and presented as geometric mean ± SD. Comparison between two groups was performed using the two-tailed Mann–Whitney U-test unless otherwise noted. Comparisons of multiple groups were performed using one-way ANOVA followed by a Tukey’s post hoc test. Comparison of survival curves was performed using a pairwise log-rank test. Comparison of viral titers was performed using lognormal one-way ANOVA followed by a Tukey’s post hoc test. P values of 0.05 or less were considered significant.

## Data availability

All data supporting the results of this study are available within the main text and its supplementary information. 3D negative-stain reconstructions for polyclonal Fab-HA complexes have been deposited in the Electron Microscopy Databank (EMDB, http://www.emdataresource.org/) and are available under the following accession codes: EMD-78950, EMD-78952, EDM-78953.

## Acknowledgments

We thank members of the Xu and Halfmann laboratories for fruitful discussions on the project. We thank P.S. Kim for helpful discussions. We thank the 3D Cell Electron Microscopy Core Facility, Cryo-Electron Microscopy Research Center, Biophysics Instrumentation Facility, and the Influenza Research Institute at the University of Wisconsin-Madison for technical support. This work was supported by startup funds from the Department of Biochemistry at University of Wisconsin-Madison to D.X. and by the University of Wisconsin’s Office of the Vice Chancellor for Research (OVCR) to P.J.H. N.A.M. was supported by the Molecular Biophysics Training Program (MBTP) at the University of Wisconsin-Madison (T32GM158463).

