## supplementary figures and legends for "Designing a reoriented H5 hemagglutinin to generate stem-directed, heterosubtypic immunity to influenza A viruses"

### Supplementary Figure and Legends

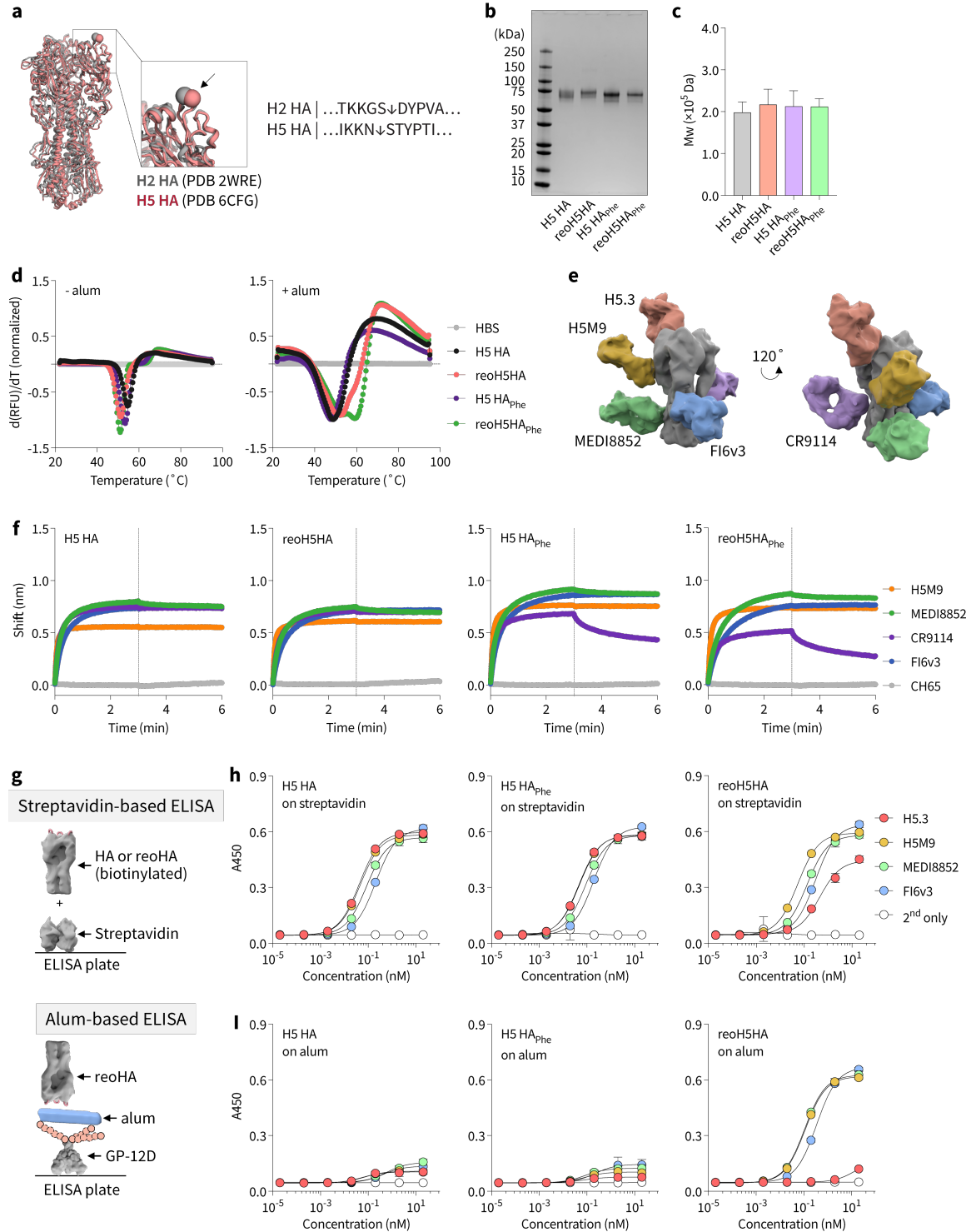

**Supplementary Figure 1. Biochemical characterization of wild-type and modified H5 HAs.**

**a**, Superposition of H2 HA (A/Japan/305/1957, PDB ID: 2WRE) and H5 HA (VN/04, PDB ID: 6CFG) allowed identification of oligoD insertion site on HA-head. Arrows indicate oligoD

insertion sites. **b**, Protein gel electrophoresis of wild-type and modified H5 HA proteins. Gel was stained with Coomassie brilliant blue. **c**, Molecular weight (Mw) calculation of wild-type and modified H5 HA proteins from the SEC-MALS analysis. Data are presented as mean ± s.d. as reported from the ASTRA software. **d**, Representative thermal melting profiles of wild-type and modified H5 HA proteins in the absence or presence of alum measured by differential scanning fluorimetry. HEPES buffer (HBS, 20 mM HEPES, 150 mM NaCl, pH 7.4) served as a negative control. **e**, Epitopes targeted by head- (H5.3), VE- (H5M9) and stem-directed mAbs (MEDI8852 and FI6v3) used to assess epitope presentation or accessibility. **f**, BLI binding profiles of wild-type and modified H5 HA proteins with HA-specific mAbs (H5M9, MEDI8852, CR9114, or FI6v3). A head-specific mAb (CH65) to H1 HA served as a negative control. Vertical dashed lines indicate the beginning of dissociation. **g**, Comparison of streptavidin- and alum-based ELISAs for analysis of antigen orientation. **h,i**, Binding of head- (H5.3), VE- or stem-directed mAbs (MEDI8852 or FI6v3) to H5 HA, H5 HA<sub>Phe</sub> or reoH5HA on **(h)** streptavidin-coated or **(i)** alum-coated ELISA plates. Data are presented as mean ± s.d. ($n = 4$ technical replicates).

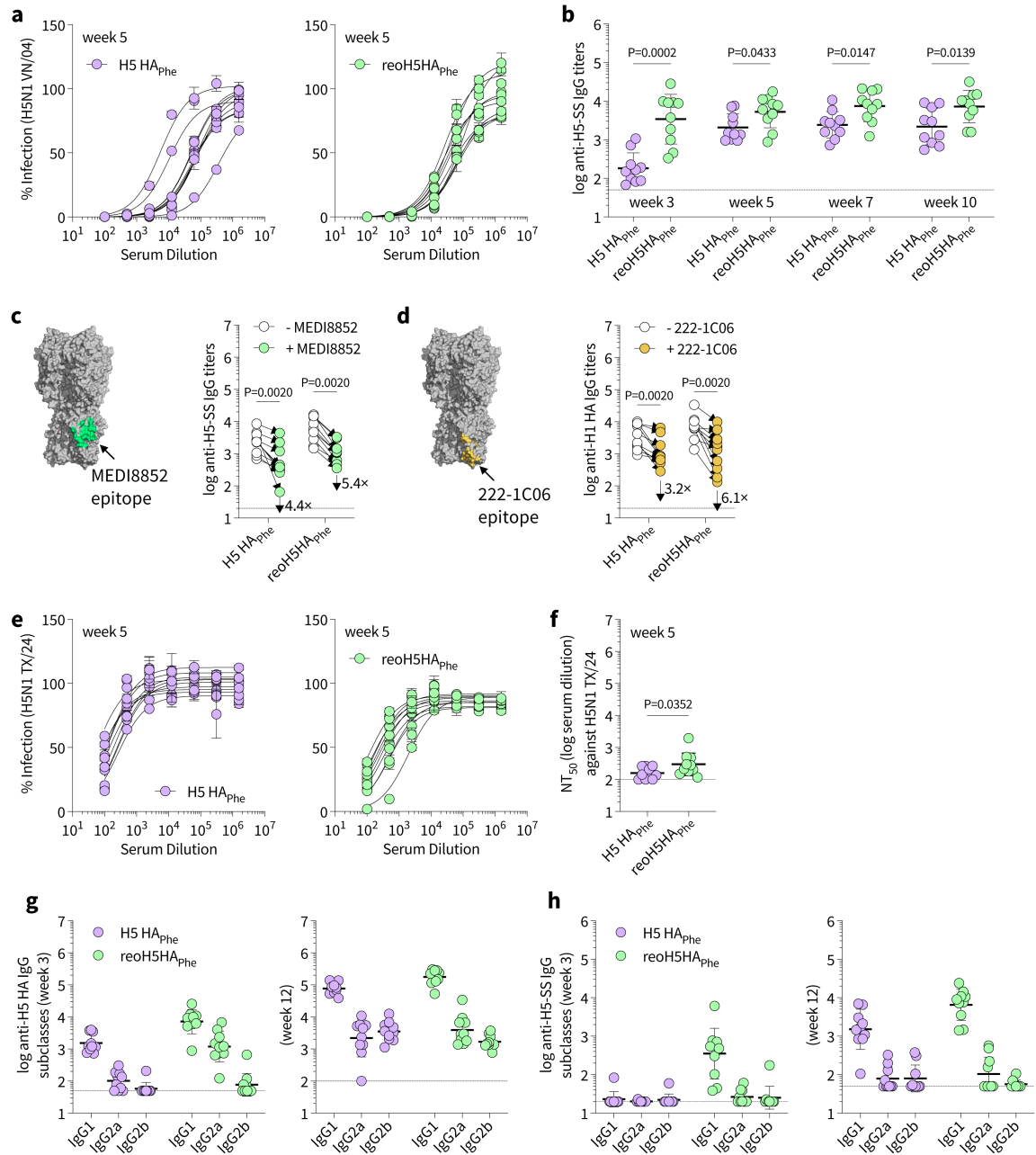

**Supplementary Figure 2. Analyses of antibody responses to the prime-boost immunization with H5 HA<sub>Phe</sub> or reoH5HA<sub>Phe</sub>.** **a**, Neutralization of the homologous H5N1 pseudovirus (VN/04) by week 5 antisera. Data are presented as mean ± s.d. ($n = 2$ technical replicates). **b**, Comparison of serum H5-SS-specific IgG titers between the two groups over time. **c,d**, Serum binding titers to H5-SS or H1 HA in the presence of a competing mAb, **(c)** MEDI8852 or **(d)** 222-1C06, respectively. Fold change in IgG titers is indicated by arrows with numbers. Comparison of two groups was performed using the Wilcoxon matched-pairs signed rank test. **e**, Cross-neutralization of a heterologous H5N1 pseudovirus (TX/24) by week 5 antisera. Data are presented as mean ±

s.d. ($n = 2$ technical replicates). **f**, Comparison of $NT_{50}$ against the heterologous H5N1 pseudovirus (TX/24). **g**, H5 HA-, and **h**, Serum H5-SS-specific IgG subclasses of week 3 or week 12 antisera. Data are presented as geometric mean ± s.d. of the $\log_{10}$-transformed values in **b-d,f-h**. Dashed lines indicate the limit of quantification in **b-d,f-h**. Comparison of two groups was performed using the two-tailed Mann-Whitney U-test in **b,f**. P values of 0.05 or less were plotted and considered significant.

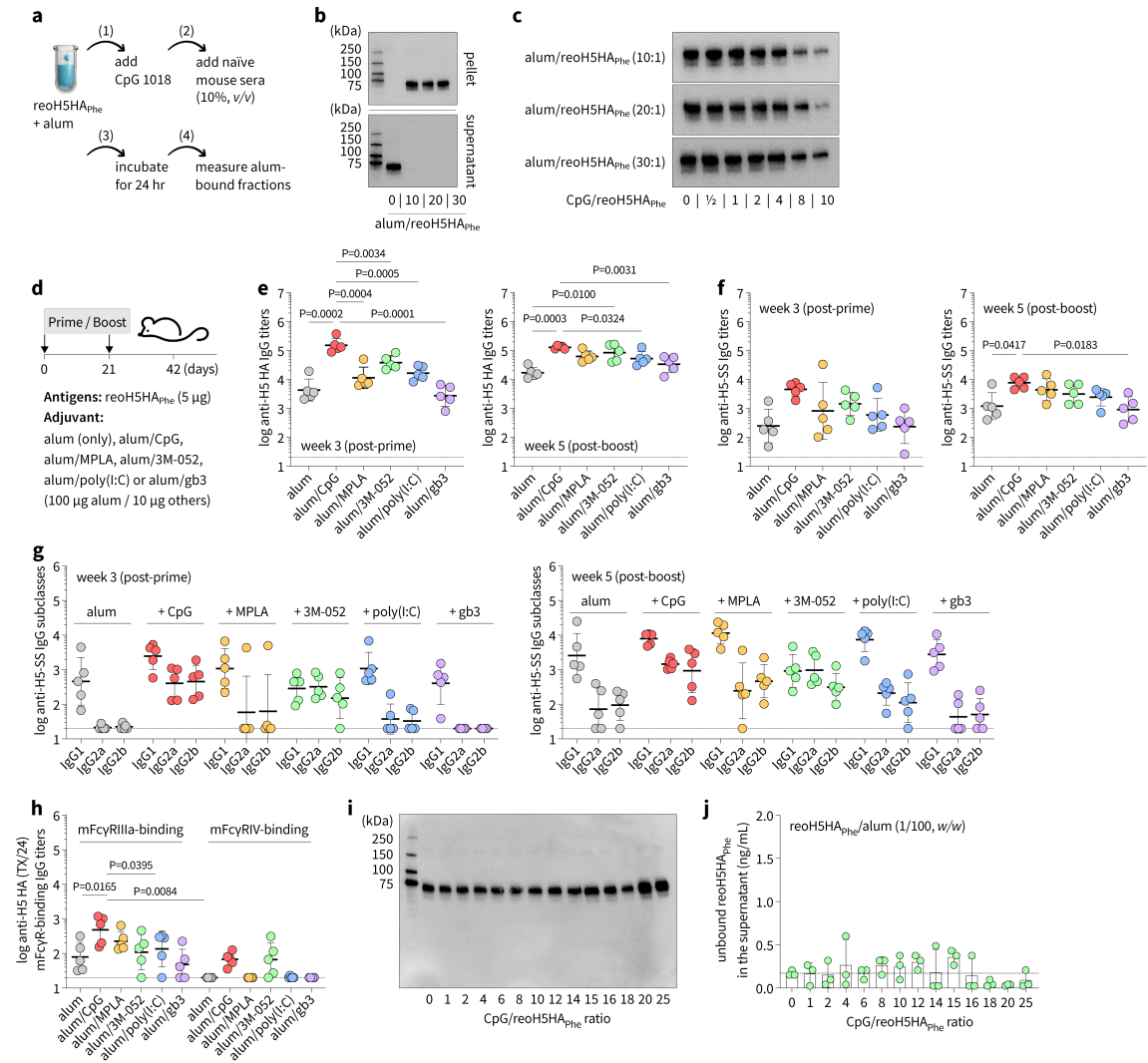

**Supplementary Figure 3. Optimizing adjuvant combinations to achieve a balanced Th1/Th2 response.** **a**, Titration of alum/CpG ratios that are compatible with reoH5HA<sub>Phe</sub>. reoH5HA<sub>Phe</sub> was pre-mixed with alum at room temperature for 30 min followed by addition of CpG. After 30 min, naïve mouse serum (10%, v/v) was added for further incubation at 37°C for 24 hr. After centrifugation, the rinsed pellet was analyzed by Western blots to detect alum-bound reoH5HA<sub>Phe</sub>. The amount of unbound reoH5HA<sub>Phe</sub> in the supernatant was quantified by Western blots or ELISA. **b,c**, Western-blot detection of alum-bound reoH5HA<sub>Phe</sub> in the (**b**) absence or (**c**) presence of CpG. **d**, A prime-boost immunization with reoH5HA<sub>Phe</sub> and different alum-based adjuvant combinations in female BALB/c mice ($n = 5$ per group). **e**, Serum H5 HA- or **f**, H5-SS-specific IgG titers post-prime or post-boost. **g**, Serum H5-SS-specific IgG subclasses post-prime or post-boost. **h**, Mouse FcγR-binding IgG titers (mFcγRIIIa or mFcγRIV) of week 5 antisera that are cross-reactive to H5 HA (TX/24). **i,j**, Titration of the permissive amount of CpG compatible

with reoH5HA<sub>Phe</sub>/alum at 1/100 (*w/w*). Western-blot analysis of alum-bound reoH5HA<sub>Phe</sub> in the pellet (**i**) and ELISA analysis of unbound reoH5HA<sub>Phe</sub> in the supernatant (**j**, *n* = 3 technical replicates). Data are presented as geometric mean ± s.d. of the log<sub>10</sub>-transformed values in **e-h** or mean ± s.d. in **j**. Dashed lines indicate the limit of quantification in **e-h,j**. Comparison of IgG titers was performed using one-way ANOVA followed by a Tukey's post-hoc test in **e,f,h**. P values of 0.05 or less were plotted and considered significant.

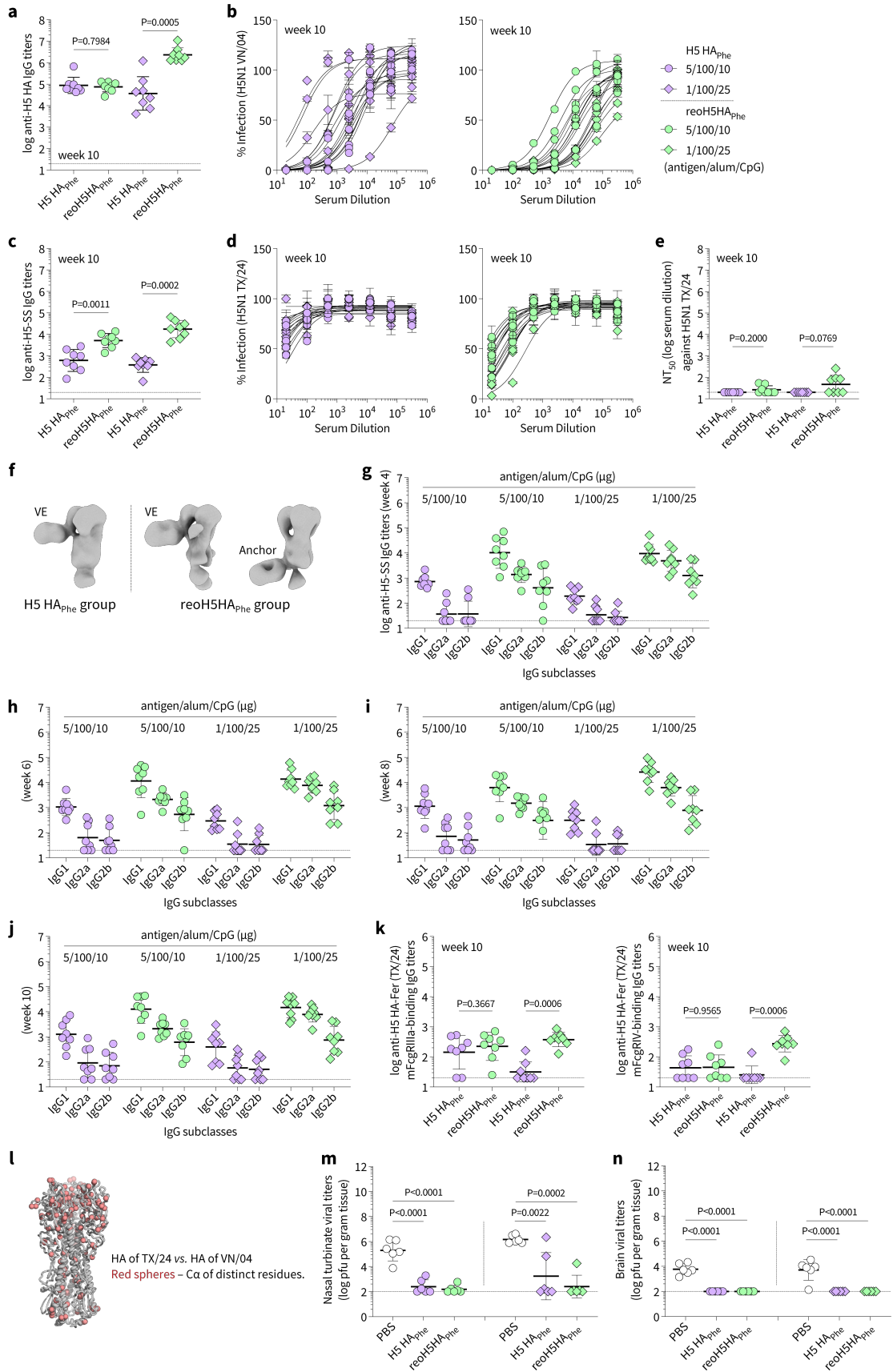

**Supplementary Figure 4. Analyses of antibody responses to the single-dose immunization with H5 HA<sub>Phe</sub> or reoH5HA<sub>Phe</sub>.** **a**, Comparison of endpoint (week 10) H5 HA-specific IgG titers. **b**, Neutralization of the homologous H5N1 pseudovirus (VN/04) by week 10 antisera. **c**, Comparison of endpoint (week 10) H5-SS-specific IgG titers. **e**, Neutralization and **e**, NT<sub>50</sub> against a heterologous H5N1 pseudovirus (TX/24) by week 10 antisera. **f**, Representative *ab-initio* reconstructions of polyclonal immune complexes with H5 HA (TX/24) for the two groups. **g-j**, Serum H5-SS-specific IgG subclasses (**g**) 4, (**h**) 6, (**i**) 8, or (**j**) 10 weeks post-injection. **k**, mFcγRIIIa- and mFcγRIV-binding IgG titers of antisera that are cross-reactive to H5 HA (TX/24). **l**, Structural mapping of amino acid differences in HA between the vaccine antigen (VN/04) and the challenge virus (TX/24). Red spheres represent Cα of distinct amino acids. **m,n**, Viral titers within (**m**) nasal turbinate and (**n**) brain tissues three days post-infection with H5N1 virus (10× mLD<sub>50</sub>, TX/24). Comparison of viral titers from multiple groups was performed using lognormal one-way ANOVA followed by a Tukey's post-hoc test. Data are presented as geometric mean ± s.d. of the log<sub>10</sub>-transformed values in **a,c,e,g-k,m,n**, or as mean ± s.d. in **b,d**. Dashed lines indicate the limit of quantification in **a,c,e,g-k,m,n**. Comparison of two groups was performed using the two-tailed Mann-Whitney U-test. P values of 0.05 or less were plotted and considered significant.

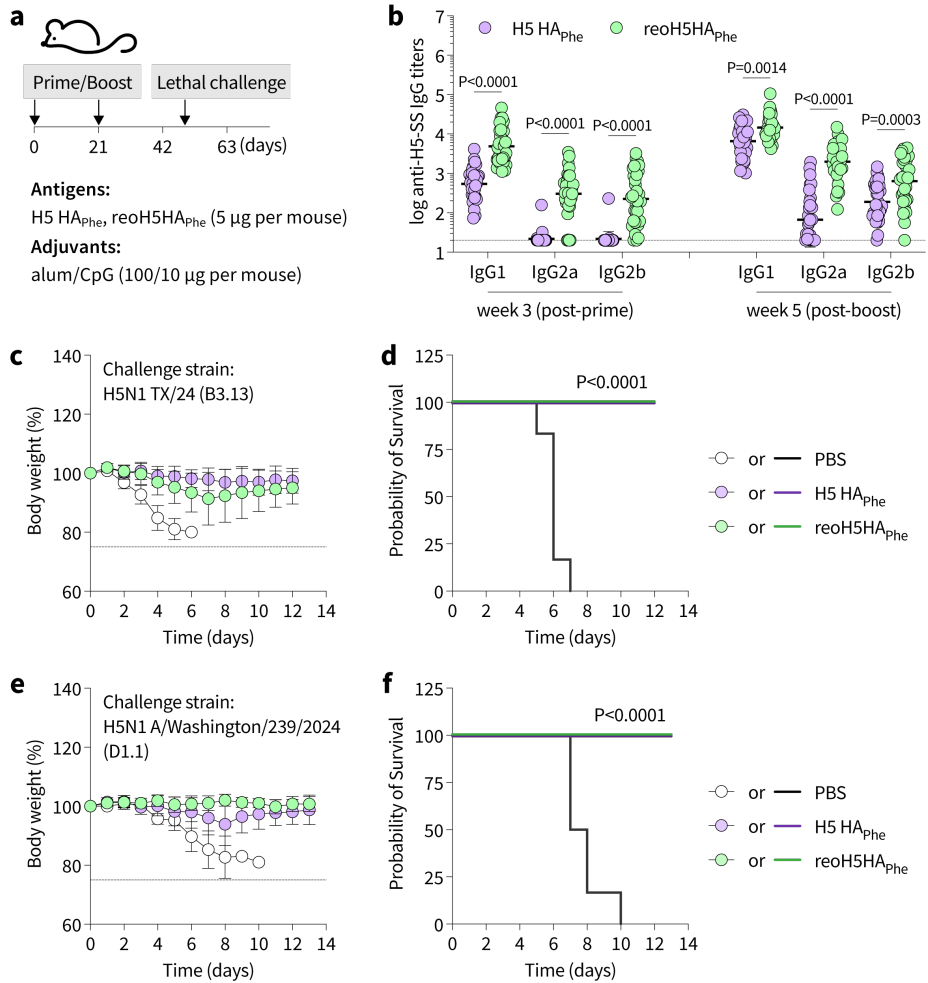

**Supplementary Figure 5. Analysis of antibody responses and protective efficacy following the prime-boost immunization with H5 HA<sub>Phe</sub> or reoH5HA<sub>Phe</sub>.** **a**, A prime-boost immunization with H5 HA<sub>Phe</sub> or reoH5HA<sub>Phe</sub> (5 µg per mouse) adjuvanted with alum/CpG (100/10 µg per mouse) *via* intramuscular injection in female BALB/c mice ($n = 32$ per group). **b**, Serum H5-SS-specific IgG subclasses post-prime or post-boost. Data are presented as geometric mean ± s.d. of the log<sub>10</sub>-transformed values. Comparison of two groups was performed using the two-tailed Mann-Whitney U-test. **c-f**, Body weight loss (**c,e**) and kaplan-meier survival curves (**d,f**) of mice after lethal challenge ($10 \times \text{mLD}_{50}$) with the HPAI H5N1 viruses with different genotypes (B3.13 genotype – TX/24 vs. D1.1 genotype – A/Washington/239/2024). Data are presented as mean ± s.d. and dashed lines indicate 25% body weight loss in **c,e**. P values of 0.05 or less were plotted and considered significant.

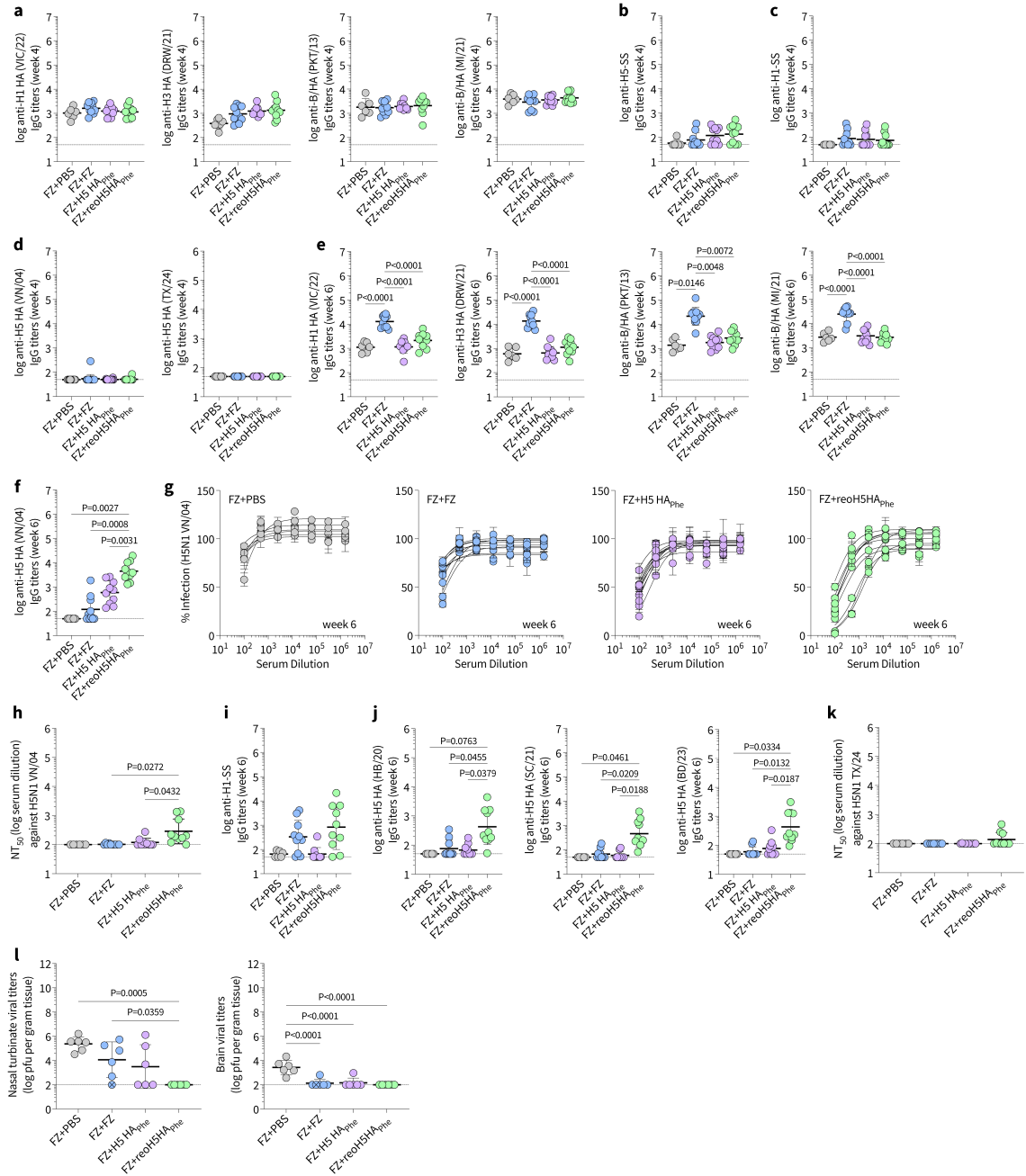

**Supplementary Figure 6. Analysis of antibody responses following the prime immunization with FZ and different booster immunizations. a-e**, Serum IgG titers against (a) HAs from FZ vaccine strains, including H1 HA (VIC/22), H3 HA (DRW/21), B/HA (PKT/13), B/HA (MI/21), (b) H5-SS, (c) H1-SS, and (d) H5 HA (VN/04 or TX/24) 4 weeks post-prime. **e,f**, Serum IgG titers against (e) HAs from FZ vaccine strains or (f) H5 HA (VN/04) 2 weeks post-boost. **g**, Neutralization and **h**, NT<sub>50</sub> against the homologous H5N1 pseudovirus (VN/04) 2 weeks post-boost. **i,j**, Serum IgG titers against (i) H1-SS, or (j) HPAI H5 HAs (HB/20, SC/21, BD/23) 2

weeks post-boost. **k**, NT<sub>50</sub> against the heterologous H5N1 pseudovirus (TX/24) 2 weeks post-boost. **l**, Viral titers within nasal turbinate and brain tissues three days post-infection with H5N1 virus (10× mLD<sub>50</sub>, TX/24). Comparison of viral titers from multiple groups was performed using lognormal one-way ANOVA followed by a Tukey's post-hoc test. Data are presented as geometric mean ± s.d. of the log<sub>10</sub>-transformed values in **a-f,h-l**, or mean ± s.d. in **g**. Dashed lines indicate the limit of quantification. Comparison of multiple groups was performed using one-way ANOVA followed by a Tukey's post-hoc test. P values of 0.05 or less were plotted and considered significant.

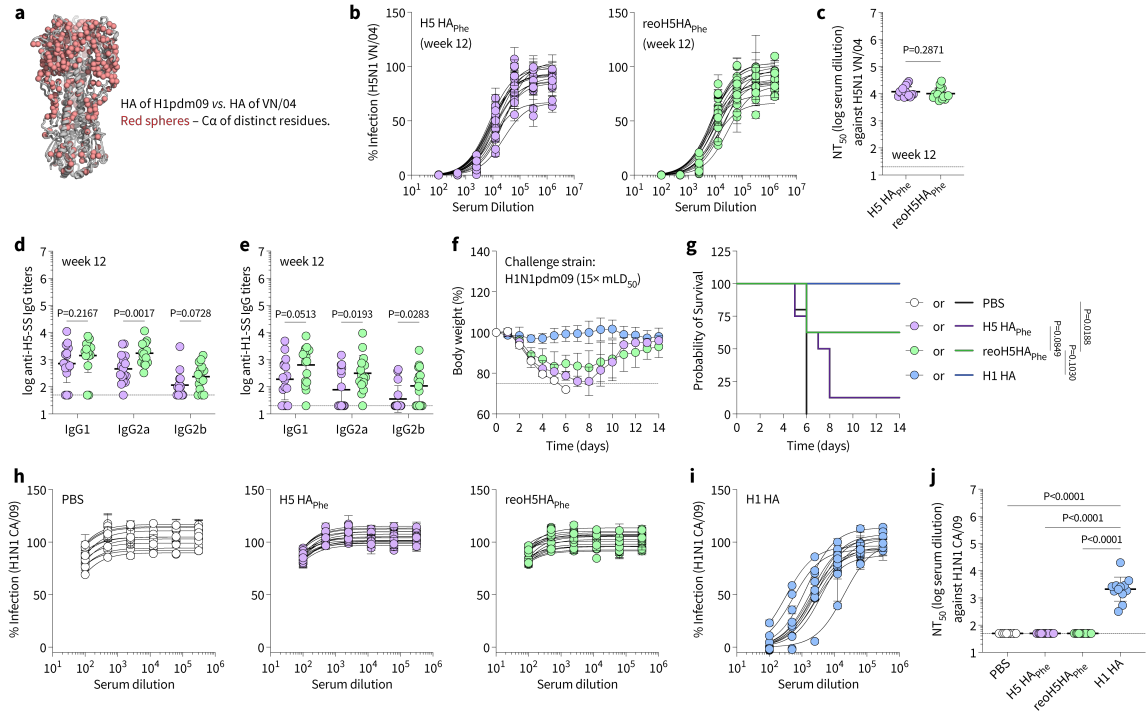

**Supplementary Figure 7. Analysis of antibody responses and protective efficacy following three immunizations with H5 HA<sub>Phe</sub> or reoH5HA<sub>Phe</sub>.** **a**, Structural mapping of amino acid differences in HA between H5 (VN/04) and H1 HAs (H1N1pdm09). Red spheres represent Ca of distinct amino acids. **b**, Neutralization and **c**, NT<sub>50</sub> against the homologous H5N1 pseudovirus (VN/04) by week 12 antisera. **d**, Serum H5-SS- and **e**, H1-SS-specific IgG subclasses by week 12 antisera. **f**, Body weight loss and **g**, Kaplan-Meier survival curves of mice after lethal challenge (15× mLD<sub>50</sub>) with H1N1pdm09. Comparison of survival curves was performed using a pairwise log-rank test. **h**, **i**, Cross-neutralization and **j**, NT<sub>50</sub> against the wild-type H1N1 virus (CA/09) by week 12 antisera. Data are presented as mean ± s.d. in **b**, **f**, **h**, **i**, or geometric mean ± s.d. of the log<sub>10</sub>-transformed values in **c**-**e**, **j**. Dashed lines indicate limit of quantification in **c**-**e**, **j** or 25% body weight loss in **f**. Comparison of two groups was performed using the two-tailed Mann-Whitney U-test. Comparison of multiple groups was performed using one-way ANOVA followed by a Tukey's post-hoc test. P values of 0.05 or less were plotted and considered significant.

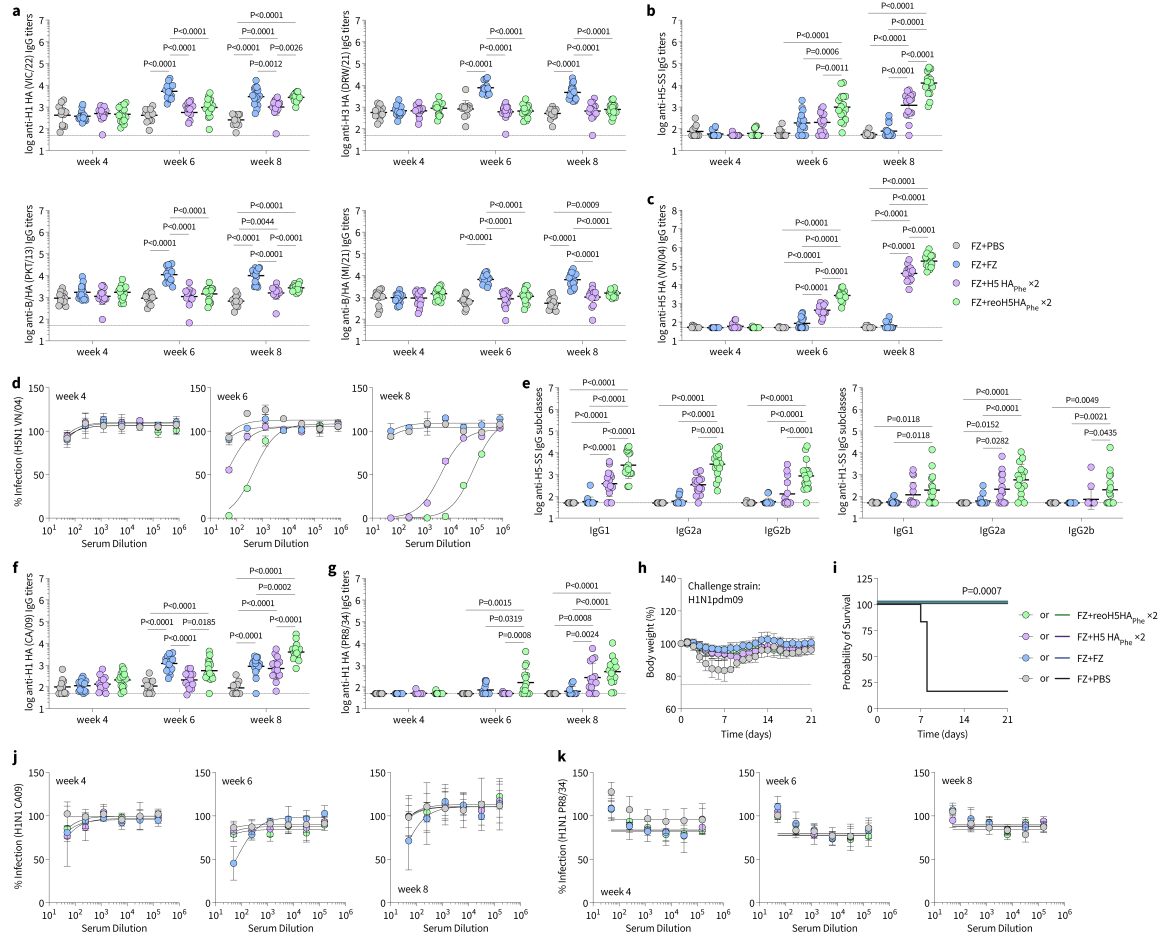

**Supplementary Figure 8. Analysis of antibody responses and heterosubtypic protection following the prime immunization with FZ and different booster immunizations. a,b,** Serum IgG titers against (a) HAs from FZ vaccine strains and (b) H5 HA (VN/04) post-prime (week 4) or post-boost (week 6 and 8). c, Neutralization of the homologous H5N1 pseudovirus (VN/04) post-prime or post-boost. Antisera were pooled from each group for the assay ($n = 4$ technical replicates). d, Serum H5-SS-specific IgG titers over time. e, Serum H5-SS- or H1-SS-specific IgG subclasses by week 8 antisera. f,g, Cross-reactive IgG titers against (f) H1 CA/09 or (g) H1 PR8/34 post-prime or post-boost. h, Body weight loss and i, Kaplan-Meier survival curves of mice after lethal challenge ($10 \times \text{mLD}_{50}$) with the heterosubtypic H1N1pdm09 virus. Comparison of survival curves was performed using a pairwise log-rank test. j,k, Neutralization of the wild-type (j) H1N1 CA/09 or (k) PR8/34 viruses post-prime or post-boost. Antisera were pooled from each group for the analysis ($n = 4$ technical replicates). Data are presented as geometric mean ± s.d. of the $\log_{10}$-transformed values in a-c,e-g or mean ± s.d. in d,h,j,k. Dashed lines indicate the limit of quantification in a-c,e-g or 25% body weight loss in h. Comparison of

multiple groups was performed using one-way ANOVA followed by a Tukey's post-hoc test. P values of 0.05 or less were plotted and considered significant.
